# A new cortical parcellation based on systematic review of primate anatomical tracing studies on corticostriatal projections

**DOI:** 10.1101/2022.06.20.496804

**Authors:** Tovy Dinh, Stener Nerland, Ivan I. Maximov, Claudia Barth, Anthony C. Vernon, Ingrid Agartz, Kjetil Nordbø Jørgensen

## Abstract

Corticostriatal projections form the input level of a circuitry that connects the cerebral cortex, basal ganglia, and thalamus. Three distinct, functional subcircuits exist according to the tripartite model: Sensorimotor cortices projecting mainly to the dorsolateral striatum; associative cortices projecting to the dorsomedial striatum and limbic cortices projecting to the ventral striatum. However, there is to date no atlas that allows researchers to label cortical projection areas belonging to each of these subcircuits separately.

To address this research gap, the aim of this study was threefold: First, to systematically review anatomical tracing studies that focused on corticostriatal projections in non-human primates, and to classify their findings according to the tripartite model. Second, to develop an atlas of the human cerebral cortex based on this classification. Third, to test the hypothesis that labels in this atlas show structural connectivity with specific striatal subregions in humans using diffusion-based tractography in a sample of 24 healthy participants.

In total, 98 studies met the inclusion criteria for our systematic review. Information about projections from the cortex to the striatum was systematically extracted by Brodmann area, and cortical areas were classified by their dominant efferent projections. Taking known homological and functional similarities and differences between non-human primate and human cortical regions into account, a new human corticostriatal projection (CSP) atlas was developed. Using human diffusion-based tractography analyses, we found that the limbic and sensorimotor atlas labels showed preferential structural connectivity with the ventral and dorsolateral striatum, respectively. However, the pattern of structural connectivity for the associative label showed the greatest degree of overlap with other labels.

We provide this new atlas as a freely available tool for neuroimaging studies, where it allows for the first-time delineation of anatomically informed regions-of-interest to study functional subcircuits within the corticostriatal circuitry. This tool will enable specific investigations of subcircuits involved in the pathogenesis of neuropsychiatric illness such as schizophrenia and bipolar disorders.

**Highlights:**

- Systematic review of anatomical projections from the cerebral cortex to the striatum in non-human primates.
- Development of a novel cortical atlas for use in neuroimaging studies focusing on the corticostriatal brain circuitry.
- Tractography in human diffusion-weighted imaging data to test if associative, limbic, and sensorimotor cortical atlas labels show preferential connectivity to regions within the striatum.

## 1 Introduction

The majority of the cerebral cortex is connected to the basal ganglia via a complex brain network; the corticostriatal circuitry. This circuitry plays a crucial role in the development and coordination of adaptive goal-directed behavior [1], and its dysfunction has been associated with a variety of neuropsychiatric and neurodegenerative disorders. For instance, dopamine dysfunction in the striatum has been repeatedly reported in schizophrenia [2–5]. Striatal degeneration is a key aspect of the neuropathology in Huntington’s disease [6, 7]. Furthermore, aberrant striatal dopamine transmission is also reported in Parkinson’s disease [8], and specific parts of this circuitry are also implicated in obsessive-compulsive disorder [9] and Tourette syndrome [10–13].

Corticostriatal projections form the input stage of a complex circuitry that connects the cortex, basal ganglia, and the thalamus, with re-entrant projections to cortical regions [14]. These projections are topographically ordered and the correspondence between cortical input regions and striatal terminal zones appears to be largely preserved across vertebrate species [15, 16]. Based on this, the corticostriatal circuitry is thought to consist of several functional subcircuits, rather than being functionally homogenous. Currently, a tripartite model describing sensorimotor, limbic, and associative subcircuits [17, 18] has emerged as important [19], although models describing four [20] or five [21] different subcircuits also exist.

According to the tripartite model, the dorsolateral striatum is considered its sensorimotor region, the dorsomedial striatum its associative region, and the ventral striatum its limbic region (Figure 1 D) [17, 22–26]. It should be noted, however, that these subcircuits may not be sharply delineated. Although they were previously thought of as highly segregated, following the influential account by Alexander, DeLong, and Strick (1986) [21], there is evidence of overlap in boundary areas and integration between these circuits at the subcortical level of the circuitry [1].

**Figures 1A, B, C and D. A.**
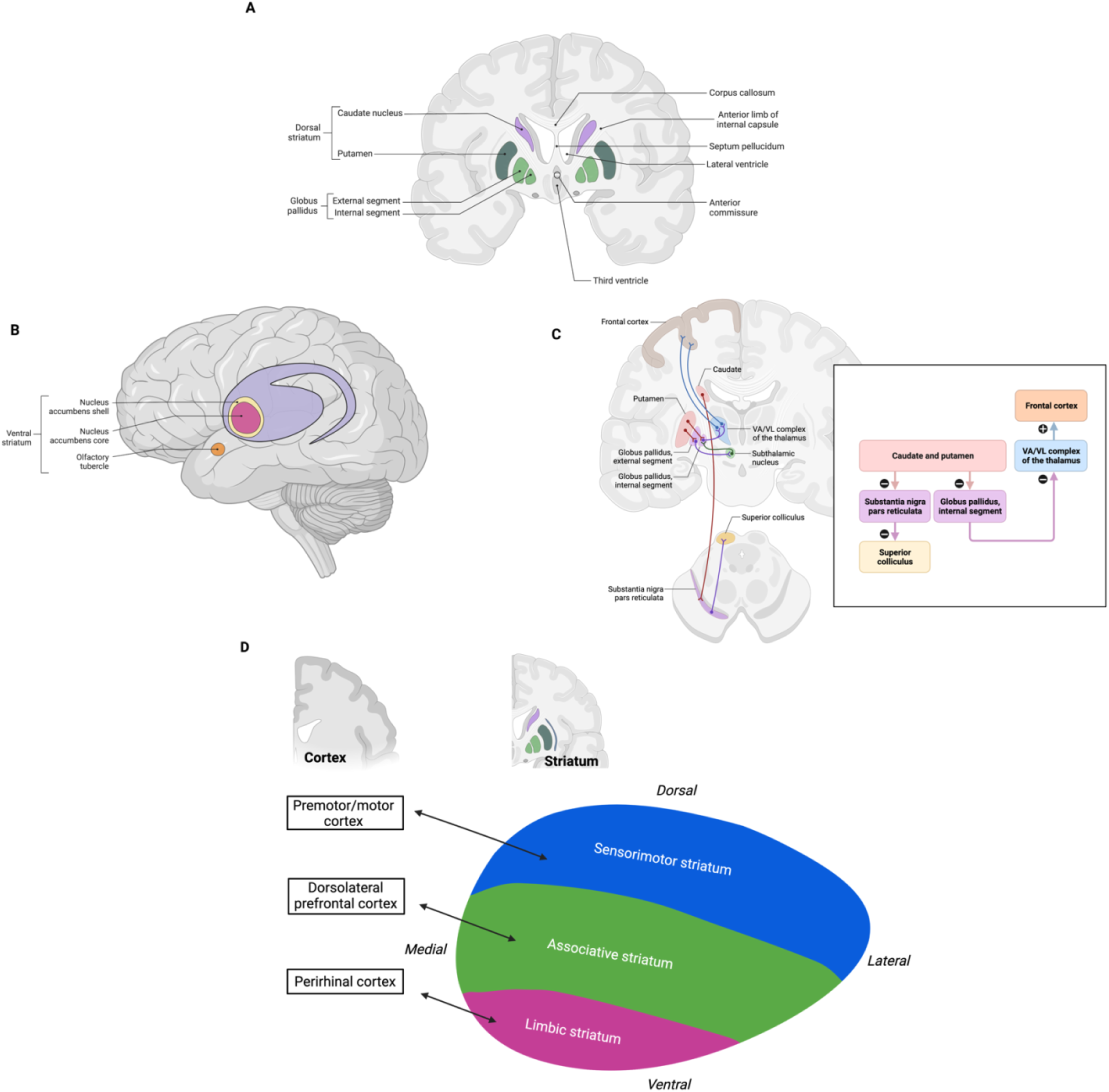
Coronal view of the anatomical position of the dorsal striatum within the brain. Adapted from “Coronal Section of the Brain (Cut at Basal Ganglia) – Dynamic Shapes”, by BioRender.com (2022). Retrieved from https://app.biorender.com/biorender-templates. **B.** Sagittal view of the anatomical position of the ventral striatum within the brain. **C.** Overview of intrinsic connectivity and outputs of the basal ganglia, here shown with the frontal cortex as an example. Reprinted from “Intrinsic Circuitry and Outputs of the Basal Ganglia”, by BioRender.com (2022). Retrieved from https://app.biorender.com/biorender-templates. **D.** Schematic topography of corticostriatal projections. The three functional subcircuits within the corticostriatal circuitry are illustrated; motor cortical areas projecting to the dorsolateral putamen, dorsolateral prefrontal cortex to the caudate and rostral putamen, and limbic areas to the ventral striatum. All created with BioRender.com. *Note: Colors should be used for all figures in print*.

This functional subdivision of the corticostriatal circuitry at both cortical and striatal levels appears to be consistent with anatomical tract-tracing studies in non-human primates [21, 27]. Furthermore, neuroimaging studies using different modalities including diffusion-weighted imaging [28], resting state functional magnetic resonance imaging (MRI) [29], positron emission tomography (PET) [30] and, in one study, combination of diffusion-weighted imaging and gene expression data [31] have corroborated this functional subdivision of the corticostriatal circuitry.

Anatomically, the primate striatum consists of the caudate nucleus and putamen, separated by the internal capsule, and the nucleus accumbens (Figure 1A). Its functional subdivision according to the tripartite model is not defined by distinct anatomical boundaries [1]. The separation between the dorsal and ventral striatum is based upon cytoarchitectonic organization: the nucleus accumbens (NAcc) is considered the main part of the *ventral striatum* and is further subdivided into its shell and core region [32], together with the olfactory tubercle [33] (Figure 1B).

Furthermore, the ventral striatum has been robustly associated with reward signaling [34, 35] and receives dopaminergic innervation primarily from the ventral tegmental area (VTA). Another cytoarchitectonic division is the organization of patch (striosome) and matrix compartments in the striatum most prominently in the head of the caudate nucleus, visible by staining for certain immunochemical markers [36–38]. The *dorsal striatum,* consisting of the caudate nucleus and putamen, receives dopaminergic innervation mainly from the substantia nigra pars compacta (SNC) [39] (Figure 1A).

The *dorsolateral striatum* (DLS) and *dorsomedial striatum* (DMS) are less distinct from each other both in terms of their cytoarchitectural organization and dopaminergic (DA) innervation. They have, however, been associated with different functional roles, wherein the DLS supports motor function, whilst the DMS supports learning as well as other cognitive functions. Corticostriatal projections regarded as sensorimotor can be identified in functional studies by stimulating sensorimotor cortical areas and probing their connectivity to the dorsolateral striatum [40–42]. If the projection targets are located in the dorsolateral striatum, the projections are considered as sensorimotor. Functionally, the DLS is found to be active during the execution of learned motor sequences, while the DMS is active during learning of novel motor sequences, pointing to the DMS being an important integrator for association of information required for the learning and execution of new motor sequences [43].

In structural and functional MRI and PET studies, it is often useful to constrain analyses to specific functional subcircuits of which corticostriatal projections are part. As an example, in schizophrenia research there is evidence of differential involvement of striatal regions. In a meta-analysis of PET studies that had investigated presynaptic dopaminergic synthesis capacity or release, McCutcheon and colleagues found evidence of dopaminergic dysfunction occurring predominantly in the associative striatum and, to a lesser extent, in the sensorimotor striatum [44]. PET studies thus provide evidence for increased dopaminergic functioning in the dorsal relative to the ventral striatum [44], and that these topographical variations of dopamine disturbances in the striatum alters symptoms in psychosis [45].

Furthermore, in a recent study topographical variations of dopamine synthesis capacity were found to relate to symptom expression in patients with first-episode psychosis, although the pattern was found for striatal regions based on functional connectivity rather than anatomical subdivision. From magnetic resonance studies, there is also evidence suggesting the involvement of corticostriatal circuits. In a diffusion-weighted imaging study, reduced frontostriatal connectivity was found in patients with chronic schizophrenia [46]. From structural magnetic resonance imaging a pattern of widespread cortical thinning has emerged [47] and specific symptom-structure relationships have been reported, such as thinning of the medial orbitofrontal cortex being associated with more negative symptoms [48, 49]. Moreover, positron emission tomography studies in schizophrenia suggest specific disturbances in dopamine transmission confined to striatal subdivisions [44, 45]. In order to test hypotheses of how such alterations may involve circuit dysfunction in specific corticostriatal subcircuits, the use of anatomical constraints is needed, not only with regards to striatal regions but also to their corresponding cortical projection areas. One approach to addressing this issue is the combination of anatomical knowledge with neuroimaging data including diffusion-weighted or functional MRI [50–52].

Comparative studies of striatal subdivisions in humans and non-human primates have shown widespread anatomical preservation of corticostriatal circuits involving all three subdivisions of the striatum [51, 53]. In a study of the reward circuit by Haber and Knutson, results from human structural and functional neuroimaging were shown to be highly correspondent to findings from primate anatomical studies [51]. These data imply a high degree of evolutionary conservation and homology of this region across primate species that would allow for the extrapolation of structural and functional features from non-human primates to humans.

In this context, in the current study, we created a new corticostriatal projection (CSP) atlas based on the anatomical tracing literature on corticostriatal projections in non-human primates. We first performed a systematic review of the existing anatomical tracing literature based on non-human primates and classified cortical projections by their striatal target zones. This classification was translated to homologous regions in the human brain, using the anatomical literature as a reference. We then created a new cortical atlas based on the PALS-B12 parcellation on the standard *fsaverage* cortical surfaces. Finally, we used probabilistic tractography of diffusion-weighted imaging (DWI) data to test the hypothesis that cortical regions of the CSP atlas would show the expected preferential connectivity to the striatum. Specifically, we expected to find preferential structural connectivity between the sensorimotor label and the dorsolateral striatum, the associative label, and the dorsomedial striatum, and between the limbic label and the ventral striatum.

## 2 Methods

### 2.1 Systematic literature search

To identify relevant studies, a MEDLINE search was conducted on September 11^th^, 2019, using the following search terms: (Cercopithecidae OR monkey OR non-human primate) AND (Cerebral Cortex OR cortex OR Limbic Lobe OR Parahippocampal Gyrus OR insula) AND (Basal Ganglia OR Corpus Striatum OR striatum OR striatal OR caudate OR putamen OR accumbens) AND (Microscopy OR Histology OR Autoradiography OR Horseradish Peroxidase OR Neural Pathways OR projections OR anterograde tracing OR retrograde tracing OR anterograde labelling OR retrograde labelling).

Medical Subject Headings (MeSH) were used where applicable, as indicated by capital letters above, and three of these search terms (Limbic lobe, Parahippocampal gyrus and Neural Pathways) were restricted using the qualifiers Anatomy & Histology or Cytology. No other search restrictions were imposed. We identified 464 articles in our search. Our search strategy is available in Supplementary materials, Section 3.

### 2.2 Study selection

We assessed the studies for relevance by manually reviewing titles, keywords, abstracts, and full text manuscripts if necessary. Studies were included if they used the following anatomical tracing techniques: anterograde and retrograde tracer injections by dye injection or autoradiographic labelling, horseradish peroxidase histochemistry, fiber degeneration and lesion studies, and if these methods were employed for tracing projections from the cerebral cortex to the striatum. We excluded studies where descriptions of striatal terminal fields were absent or imprecise. Only original reports from empirical studies were selected for review, i.e., secondary literature was not included. In total, the primary selection included 84 articles. Bibliographies of all articles were subsequently reviewed to identify additional anatomical studies on cortical projections to the striatum, leading to the inclusion of 14 more studies. Thus, our final selection comprised a total of 98 articles.

### 2.3 Strategy for extraction of information and review

#### 2.3.1 Definitions of anatomical and functional striatal regions

To compile anatomical findings about specific neuronal projections from the cerebral cortex to the striatum, we reviewed the full text of the 98 articles eligible for inclusion. First, basic information about the included studies is provided in Table 1: author(s), title and publication year, species, and study method. Second, descriptions of cortical region of origin and striatal terminal field were reviewed in detail together with figures, and included in Tables 2a-e. Third, we classified each projection originating from each area of the cerebral cortex based on their striatal termination field(s) as part of the dorsolateral, the dorsomedial and central, or the ventral striatum.

**Table 1.** Overview over included studies with author, date, primate species, investigation technique and sample size. Abbreviations for methods: AT – Anterograde tracing, ART – Anterograde radioactive tracing, EPM – Electrophysiological mapping, HRPT – Horseradish peroxidase tracing, LFD – Lesion and fiber degeneration, RT – Retrograde tracing and SDL – Silver degeneration lesion. Abbreviations for primate species: AG - Ateles geoffroyi, AT – Aotus trivirgatus, C – Cebus species, CAeP - Cercopithecus aethiops pygeryhtrus, CAl – Cebus albifrons, CAp – Cebus apella, HL – Hylobates lar, M – Macaque species, MA - Macaca arctoides, MFa – Macaca fascicularis, MFu – Macaca fuscata, MM – Macaca mulatta, MN – Macaca nemestrina, MR – Macaca radiata, PT – Pan troglodytes, SF – Saguinus fuscicollis, SO - Saguinus oedipus, and SS – Saimiri sciureus.

| Author name. | Year | Species | Method | Sample size |  | Author name | Year | Species | Method | Sample size |
| --- | --- | --- | --- | --- | --- | --- | --- | --- | --- | --- |
| Alipour et al. [66] | 2002 | SF | AT with EPM | 3 |  | Griggs et al. [67] | 2017 | MM | RT with EPM | 2 |
| Arikuni & Kubota [68] | 1986 | MM/MFu | RT | 3 / 3 |  | Haber et al. [69] | 1995 | MM/MN | AT with EPM | 8 / 14 |
| Astruc [70] | 1971 | MM | LFD | 8 |  | Haber et al. [71] | 2006 | MN/MFa | AT | 10 / 2 |
| Averbeck et al. [72] | 2014 | MN/MFa | AT and RT | 20 / 8 |  | Hatanaka et al. [73] | 2005 | MFu | AT with EPM | 7 |
| Baizer et al. [74] | 1993 | MM | AT and RT | 6 |  | Huerta & Kaas [75] | 1990 | MFa | AT with EPM | 4 |
| Baleydier & Maugiere [76] | 1980 | Not described | ART and HRPT | 13 |  | Inase et al. [77] | 1996 | MFu | AT | 4 |
| Borra et al. [78] | 2015 | MM/MFa | AT | 5 / 2 |  | Inase et al. [79] | 1999 | MFu | AT with EPM | 6 |
| Boussaoud et al. [80] | 1992 | MFa | AT and RT | 7 |  | Innocenti et al. [81] | 2017 | MFa | AT | 3 |
| Calzavara et al. [82] | 2007 | MN/MFa /MM | AT and RT | 15 / 4 / 1 |  | Johnson et al. [83] | 1968 | MM | SDL | Not described |
| Campos-Ortega et al. [84] | 1970 | MM | LFD | 21 |  | Jones et al. [85] | 1977 | SS/MM/MFa | RT | 13 / 6 / 14 |
| Choi et al. [86] | 2017a | MN/MFa/MM | AT and RT | 3 / 8 / 7 |  | Jung & Hong [87] | 2003a | MN | RT | 7 |
| Choi et al. [88] | 2017b | MN/MFa/MM | AT and RT | 3 / 4 / 3 |  | Jung & Hong [87] | 2003b | MN | RT | 7 |
| Cavada & Goldman-Rakic [89] | 1991 | MM | AT with EPM | 5 |  | Jürgens [90] | 1976 | SS | AT with EPM | 5 |
| Cheng et al. [91] | 1997 | MFu | AT | 4 |  | Jürgens [92] | 1984 | SS | AT | 6 |
| Chiba et al. [93] | 2001 | MFu | AT | 10 |  | Jürgens & Müller-Preuss [94] | 1977 | SS | AT with EPM | 6 |
| Chikama et al. [95] | 1997 | MN/MM | AT and RT | 11 |  | Kemp & Powell [96] | 1970 | M | LFD | 40 |
| Cui et al. [97] | 2003 | Cap | AT with EPM | 4 |  | Kunishio & Haber [98] | 1994 | MM | RT | 7 |
| De Vito & Smith [99] | 1964 | MN | LFD | 5 |  | Künzle [100] | 1975 | MFa | ART | 7 |
| Eblen & Graybiel [101] | 1995 | MFa/MM | AT | 12 / 1 |  | Künzle [102] | 1977 | MFa | ART | 6 |
| Ferry et al. [103] | 2000 | MFa/MN | AT and RT with EPM | 16 / 1 |  | Künzle [104] | 1978 | MFa | ART | 9 |
| Flaherty & Graybiel [105] | 1991 | SS | AT with EPM | 16 |  | Künzle & Akert [106] | 1977 | MFa | ART | 4 |
| Flaherty & Graybiel [107] | 1993 | SS | AT | 11 |  | Lapper et al. [108] | 1992 | SS | AT | 2 |
| Flaherty & Graybiel [109] | 1994 | SS | AT and RT with EPM | 14 |  | Leichnetz [110] | 1981 | MM/MFa/CAI/CAP | AT | 5 |
| Forbes & Moskowitz [111] | 1974 | SS | SDL | 29 |  | Leichnetz [112] | 1986 | MFa/MM | AT and RT | 1 / 2 |
| Freedman et al. [113] | 2000 | MM | AT | 3 |  | Leichnetz [114] | 2001 | CAP/MFa | AT and RT | 1 / 2 |
| Friedman et al. [115] | 2002 | MFa/MM | AT and RT | 18 / 10 |  | Leichnetz & Astruc [116] | 1975 | SO | SDL | 4 |
| Fudge et al. [117] | 2005 | MN/MFa | RT | 8 |  | Leichnetz & Astruc [118] | 1976 | SS | LFD | 6 |
| Gattass et al. [119] | 2014 | MM | AT and RT | 9 |  | Liles & Updyke [120] | 1985 | MM/MFa | AT with EPM | 1 / 1 |
| Gerbella et al. [121] | 2016 | MN/MFa/MM | AT | 6 / 2 / 2 |  | Maunsell & Van Essen [122] | 1983 | MFa | AT | 3 |
| Graham et al. [123] | 1979 | AT | AT | Not described |  | McFarland & Haber [124] | 2000 | MM/MN | RT | 8 |
| Goldman & Nauta [125] | 1977 | MM | AT | 2 |  | McGuire et al. [126] | 1991 | MFa | AT | 4 |
| Author name | Year | Species | Method | Sample size |  | Author name | Year | Species | Method | Sample size |
| Miyata & Sasaki [127] | 1984 | MFu | AT | 5 |  | Tokuno et al. [128] | 1999 | MFu | AT with EPM | 3 |
| Müller-Preuss & Jürgens [129] | 1976 | SS | AT with EPM | 5 |  | Ungerleider et al. [130] | 1984 | MFa | AT with EPM | 5 |
| Nakano et al. [131] | 1999 | M | AT | Not described |  | Van Hoesen et al. [132] | 1981 | MM | AT | 16 |
| Norden et al. [133] | 1978 | AT | LFD and AT | 6 |  | Weber & Yin [134] | 1984 | Macaca arctoides | AT | Not described |
| Parent & Parent [42] | 2006 | MFa | Electrophysiologically-guided single-axon AT | 9 |  | Wall et al. [135] | 1951 | MM | LFD | 3 |
| Parthasarathy et al. [136] | 1992 | MM/MFa | ART with EPM | 2 / 6 |  | Webster et al. [137] | 1993 | MM | AT | 6 |
| Parvizi et al. [138] | 2006 | MFa | AT | 8 |  | Webster et al. [137] | 1995 | MM | AT | 12 (6 adults) |
|  |  |  |  |  |  |  |  |  |  | and 6 infants) |
| Petras [139] | 1969 | AG, C, CAeP, MM, HL and PT | LFD | Not described |  | Weller et al. [140] | 2002 | AT / SS / MFa / Macaca radiata | AT | 5 / 6 / 1 / 1 |
| Petras [141] | 1971 | MM | LFD | 10 |  | Whitlock & Nauta [142] | 1956 | MM | LFD | 8 |
| Saint-Cyr et al. [143] | 1990 | MM | AT and RT with EPM | 10 |  | Whitworth et al. [144] | 1991 | AT | AT with EPM | 5 |
| Selemon & Goldman-Rakic [22] | 1985 | Not described | AT | 18 |  |  |  |  |  |  |
| Selemon & Goldman-Rakic [145] | 1988 | MM | AT | 3 |  | Yeterian & Pandya [146] | 1993 | MM | ART | 22 |
| Shook et al. [41] | 1991 | MN / Macaca arctoides | ART with EPM | 5 / 1 |  | Yeterian & Pandya [147] | 1994 | MM | RT | 3 |
| Simonyan & Jürgens [148] | 2003 | MM | AT with EPM | 3 |  | Yeterian & Pandya [149] | 1995 | MM | ART | 18 |
| Stanton et al. [150] | 1988 | MM/ MFa | AT | 9 / 1 |  | Yeterian & Pandya [151] | 1998 | MM | ART | 15 |
| Steele & Weller [152] | 1993 | SS | AT | 6 |  | Yeterian & Van Hoesen [153] | 1978 | MM | ART | 14 |
| Tachibana et al. [154] | 2004 | MFu | AT with EPM | 4 |  | Yeterian & Pandya [155] | 1991 | MM | ART | 19 |
| Takada et al. [156] | 1998a | MFu | AT with EPM | 2 |  |  |  |  |  |  |
| Takada et al. [40] | 1998b | MFu | AT with EPM | 12 |  |  |  |  |  |  |
| Takada et al. [157] | 2001 | MFu | AT with EPM | 7 |  |  |  |  |  |  |

Lastly, striatal projections of the cortical areas were then classified, as either sensorimotor if the striatal terminal field was mainly dorsolateral, associative if the striatal terminal field was mainly dorsomedial and central, or limbic if the striatal terminal field was mainly ventral.

#### 2.3.2 Classification of cortical areas by projections to striatal regions

Projections converging in or near the ventral striatum, including the nucleus accumbens, were considered as limbic. Convergent projections to the dorsolateral striatum were considered as sensorimotor, and lastly projections to the dorsomedial and central striatal region considered as associative. Thus, terminal fields and projections were classified as belonging to either the dorsolateral/sensorimotor, dorsomedial-central/associative or ventral/limbic striatal regions using several established diagrams [1, 54, 55]. As there is no clear cytoarchitectural or anatomical boundaries between striatal subregions, we relied on careful comparison of author descriptions and figures to improve our manual classification.

We considered the cortical region to be part of more than one functional circuit if this was sufficiently demonstrated by comparable studies. A manual threshold when considering overlapping regions was decided: if the number of studies showing projections to several domains was greater or equal to half the number of studies showing projections to one domain, the cortical area was considered as having projections to both domains.

To resolve conflicting information from studies when designating each region as part of the tripartite division, the number of studies and study quality was considered. Anterograde tracer studies confirmed by retrograde injections were given the highest priority, followed by anterograde tracing, and lastly lesion and other studies.

#### 2.3.3 Anatomical definitions of cortical regions in the human brain

We subsequently conducted another review to translate findings from non-human primate anatomical data to humans (Supplementary materials section 2). Among the 98 included studies from our initial literature review, we identified variation in the primate species studied. As such the cortical areas of origin were thus described using the closest related atlas to each study for the species, such as Brodmann’s [56], Von Bonin & Bailey’s [57] or Walker’s classifications [58]. By comparison of topological and cytoarchitectural homology, the cortical region was classified as the closest related Brodmann area in the primate species, after which homology with the human brain was described after assessing cytoarchitectural similarities.

Brodmann, however, did not describe certain cortical regions in detail. For these areas, we consider individual author descriptions of injection areas. Examples of areas not sufficiently described by Brodmann include area 29 [59] and parietal areas. In these areas, other cortical parcellation schemes such as Von Bonin and Bailey’s map of the macaque monkey cortex [57] were used as a supplement to identify the human homologue cortical region. Assessment of homology between primate and human brain anatomy, and primate species from which atlas regions were described, are available in Supplementary materials, section 2.

#### 2.3.4 Creation of the corticostriatal projection atlas

To construct the corticostriatal projection (CSP) atlas, we categorized regions of the previously published PALS-B12 atlas [60] based on our classification described above (Sections 2.3.2 and 2.3.3). Specifically, nine categories were used: 1) Sensorimotor, 2) Limbic, 3) Associative, 4) Sensorimotor and associative, 5) Limbic and associative, 6) Sensorimotor and limbic, 7) Sensorimotor, limbic, and associative, 8) No projection identified, 9) No homologue region (Table 2a-e). The PALS-B12 atlas is based on the Brodmann classification of cortical regions and available in FreeSurfer fsaverage space. Finally, we merged the labels together using the *mri_mergelabels* script provided in FreeSurfer version 6.0.0.

**Tables 2a-e.**
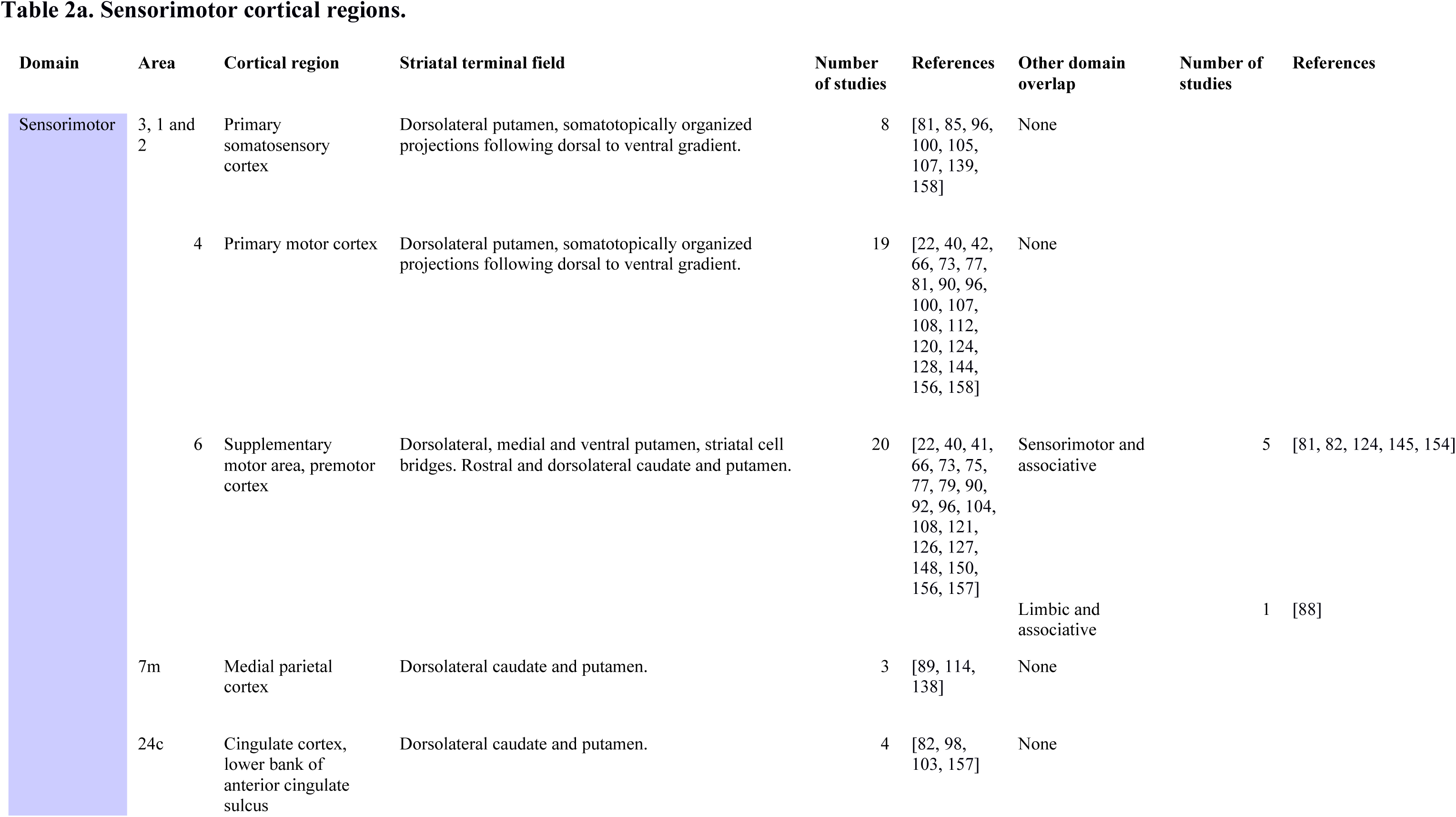
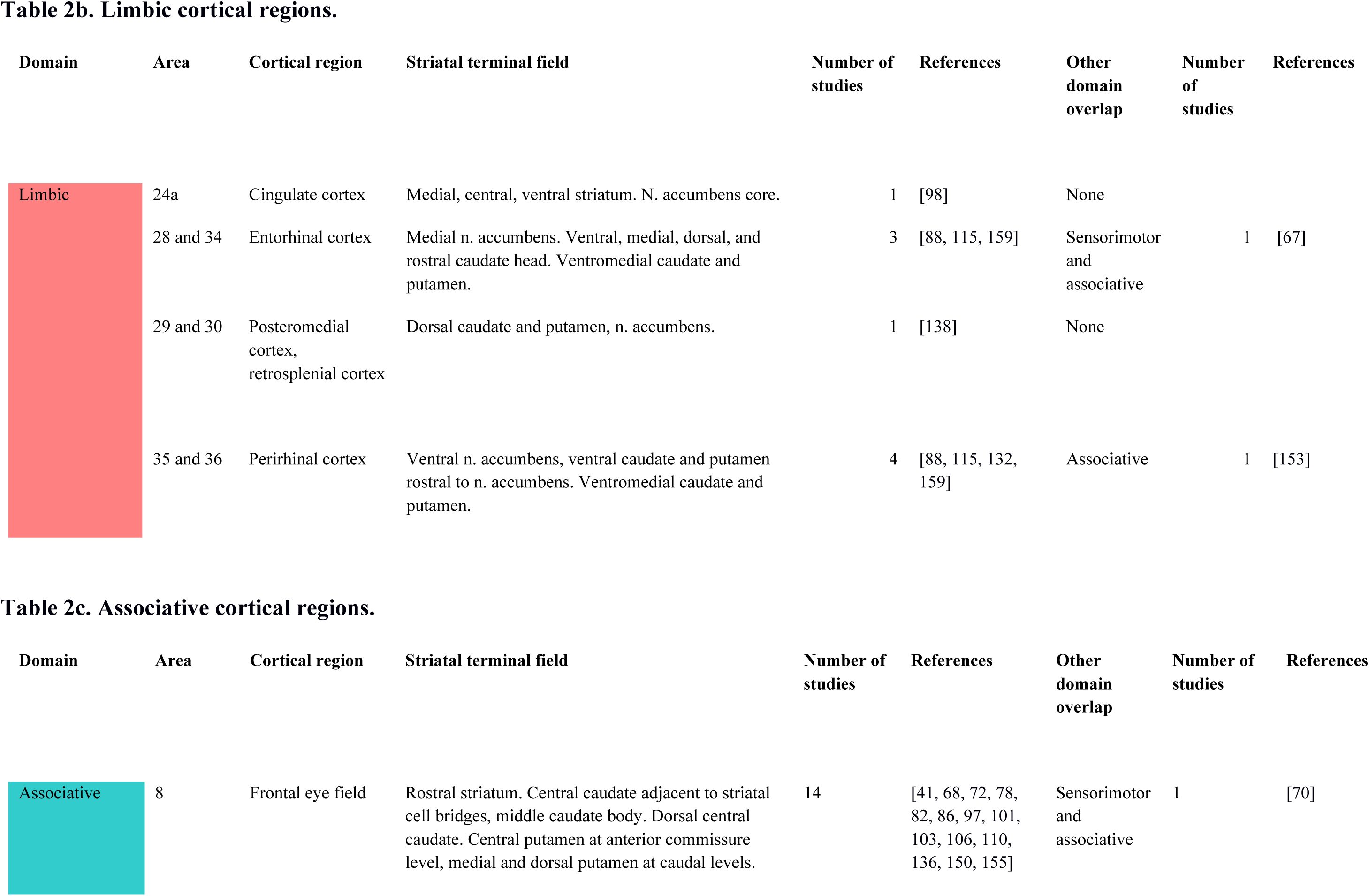
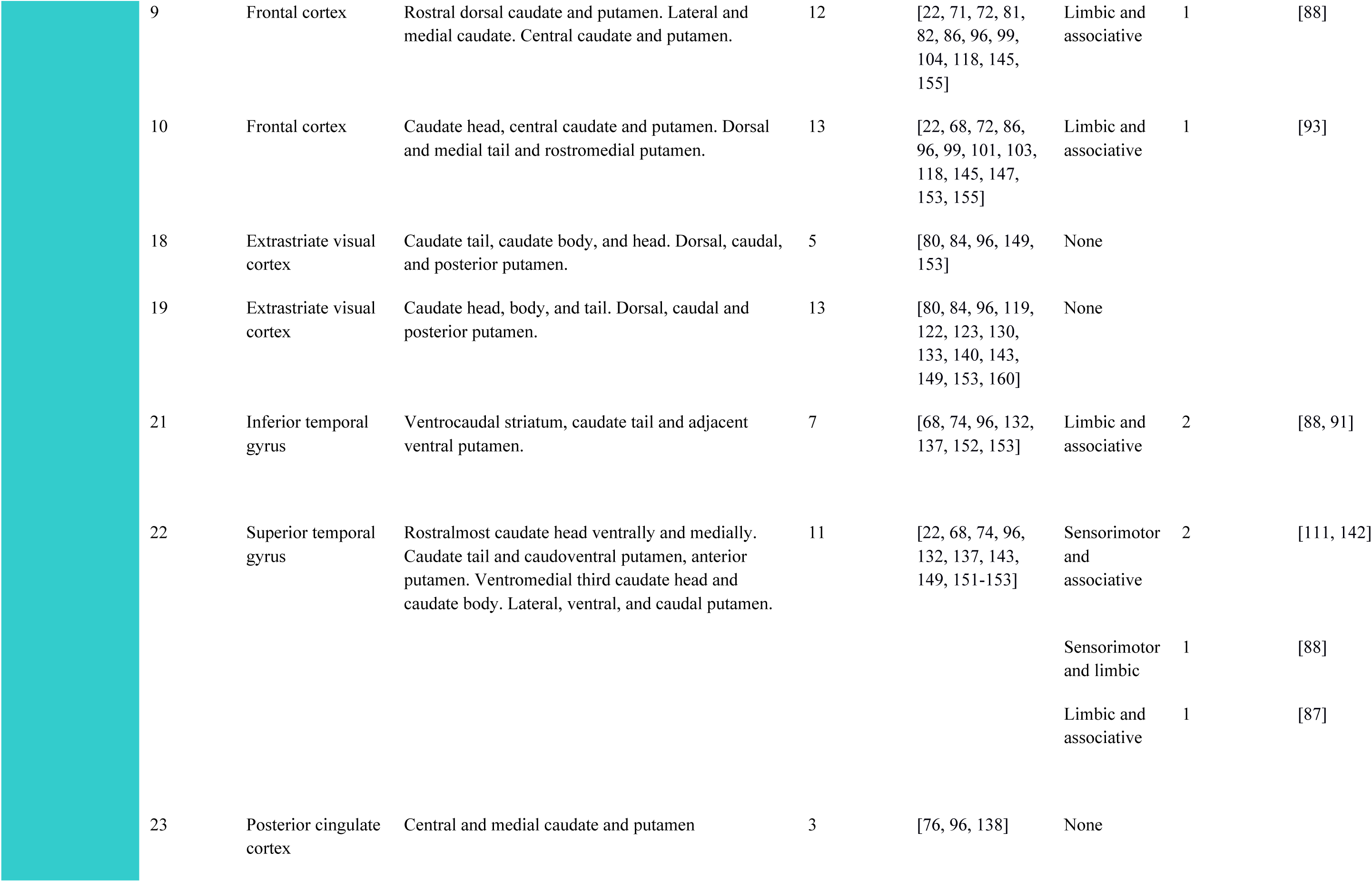
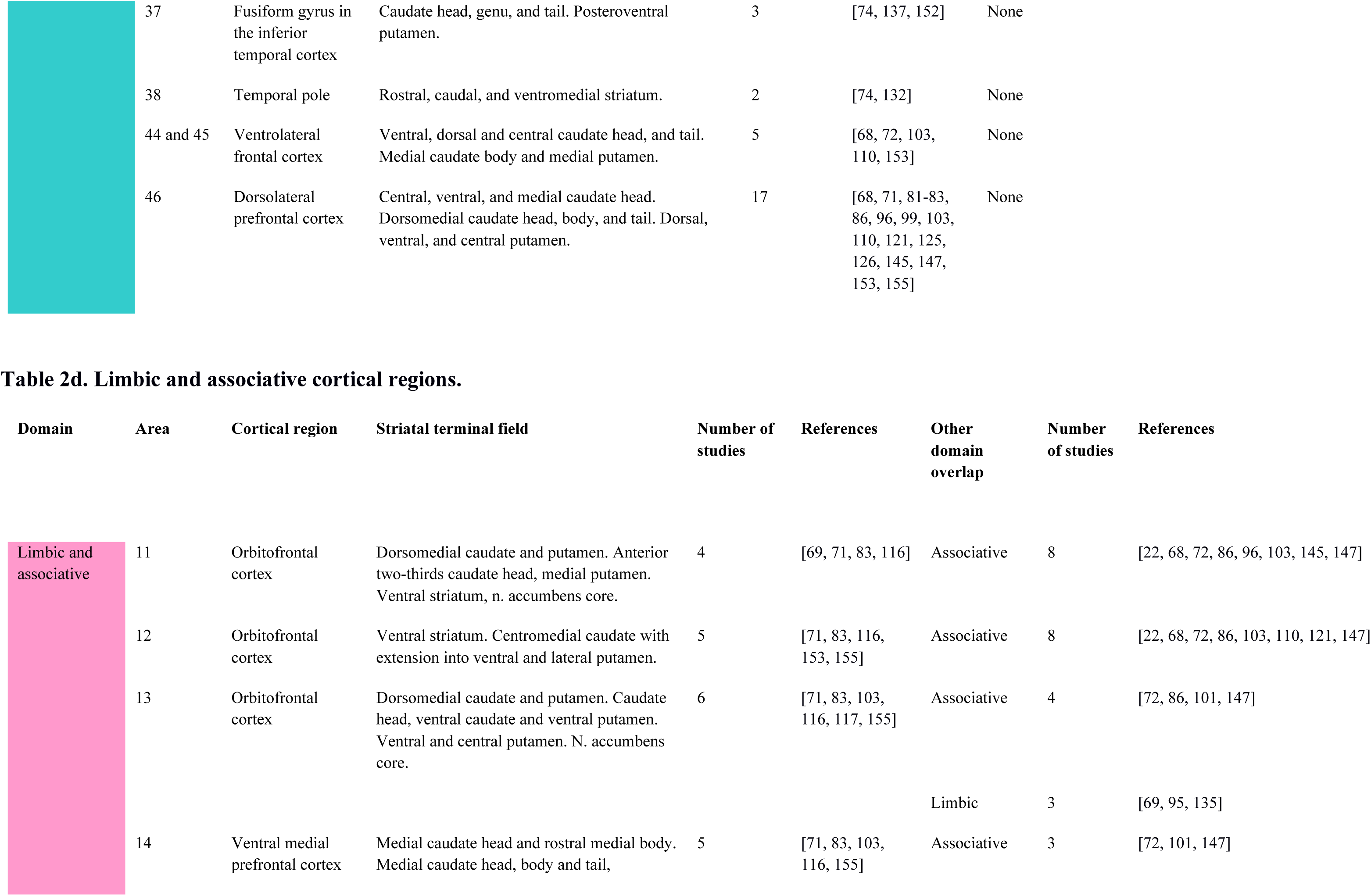
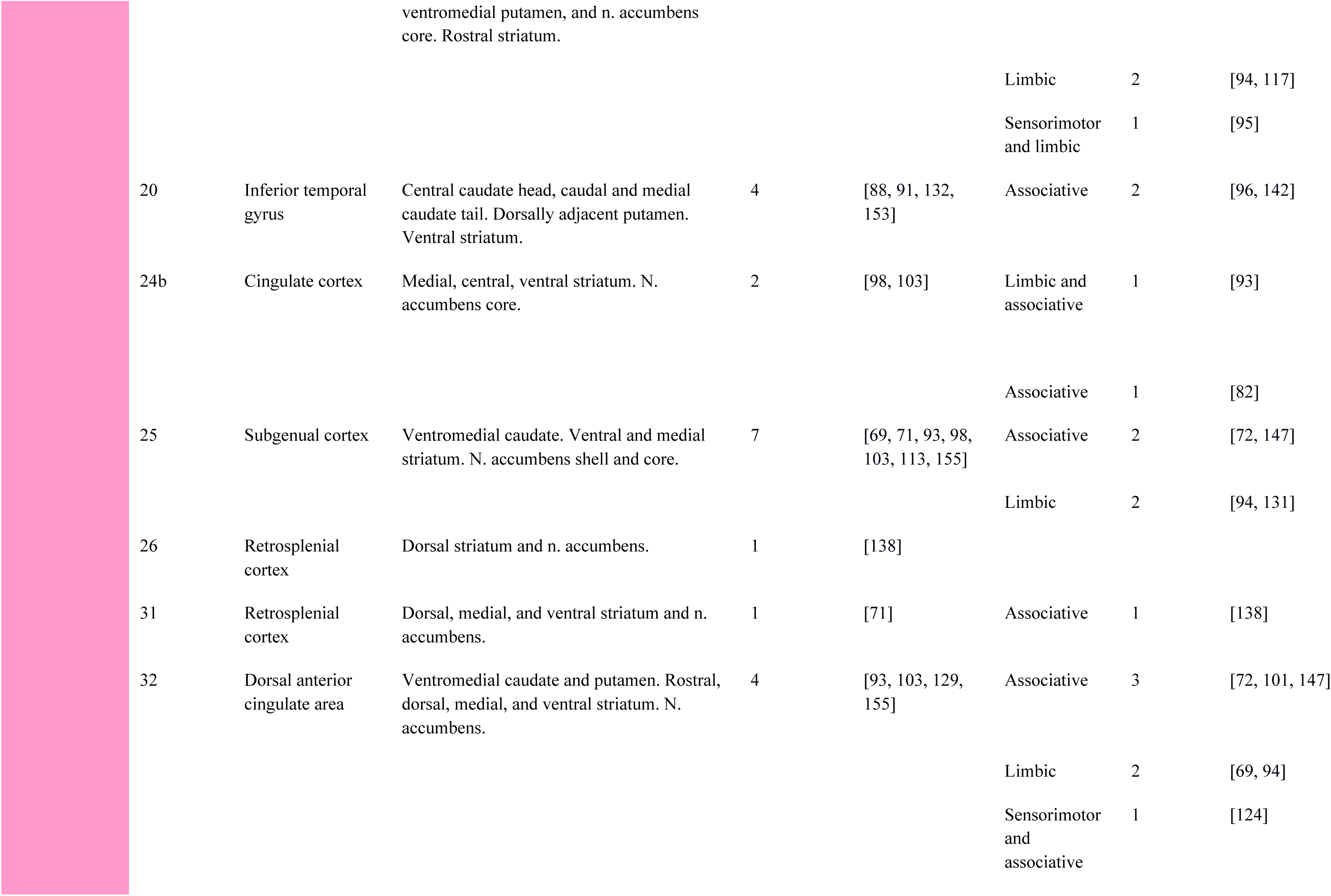
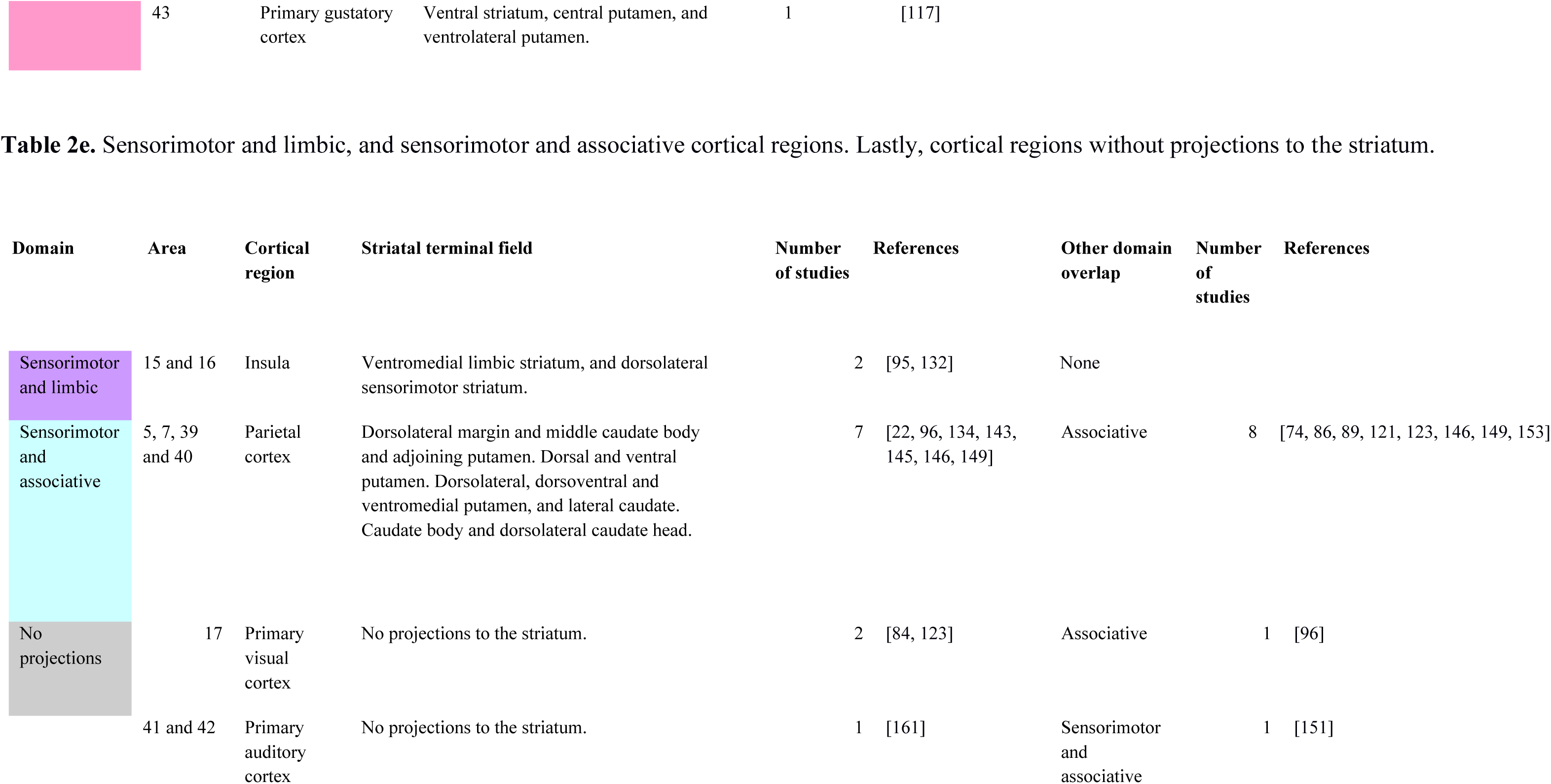
Overview of all corticostriatal projections, including corticostriatal domain, Brodmann area, cortical region, striatal terminal field, references, and domain overlaps with additional references.

#### 2.3.5 DWI-based probabilistic tractography validation

To test the hypothesis that the sensorimotor, associative, and limbic cortical labels show preferential connectivity with the dorsolateral, dorsomedial-central, and ventral striatum, respectively, diffusion-based tractography was performed. In a sample of 24 healthy participants (58.3% female, mean age 35.3 years [range: 20.1 – 51.5 years]), multi-shell diffusion weighted imaging (DWI) data was acquired at Oslo University Hospital using a GE Discovery MR750 3T scanner equipped with a 32-channel head coil. An optimized pipeline [61] was used to preprocess the diffusion-weighted data. For more details on the acquisition parameters and preprocessing steps, see the Supplementary Materials, Sections 1.1-1.2.

Probabilistic tractography was performed to estimate the voxel-wise connectivity between regions within the striatum and the cortical labels in the atlas (Section 2.3.4). In brief, data were prepared using bedpostx [62] before voxel-by-ROI connectivity estimation was performed using probtrackx2 [63], both part of the FSL toolbox version 6.0.5 [64]. Striatal seed voxels were defined by a whole striatum mask extracted from the Oxford-GSK-Imanova structural atlas [65] and cortical labels were used as regions of interest after co-registration to MNI space. Group averages for voxel-by-ROI connectivity were created in MNI space and visualized. For further details, we refer the reader to the Supplement, Sections 1.3-1.5.

#### 2.3.6 Ethics statement

Human neuroimaging data analyzed in our study were from a previous study approved by the Regional Committees for Medical Research Ethics in Southeast Norway. All participants provided written informed consent. The study was carried out in accordance with the Helsinki Declaration, and the data was handled according to guidelines set forth by the Norwegian Data Protection Authority in compliance with the European General Data Protection Regulation.

## 3. Results

### 3.1 Systematic review of the primate anatomical tract-tracing literature

Information about corticostriatal projections synthesized from this review is summarized in Tables 2a-e. Cortical projection areas are grouped by their classification into separate or overlapping functional circuits, and summary information about striatal terminal fields are described.

#### 3.1.1 Sensorimotor regions

Consistent projections were reported from Brodmann areas 3, 1, 2 and 4 to the dorsolateral putamen (Table 2a). Similarly, for area 6, 21 out of 27 studies provided evidence for (77.78%) consistent projections to the dorsolateral striatum, although projections to the dorsolateral, caudal, and ventral dorsal caudate were also found in six studies (22.22%) [81]. One retrograde study found projections from area 6 to the ventral striatum, however, here a large portion of the dorsolateral ventral striatum was injected with retrograde tracer [88]. All studies included provided evidence for projections to the dorsolateral striatum for area 7m (medial area) (Table 2a). All studies included for area 24c provided evidence for consistent projections to the dorsolateral striatum (Table 2a).

#### 3.1.2 Limbic regions

Three out of four studies (75%) provided evidence for consistent projections to the ventral striatum for areas 28 and 34 (Table 2b); except for a retrograde study with tracer injected in the caudate tail [67]. One study on areas 29 and 30 provided evidence for projections solely to the ventral striatum in one study [138]. 80% of all included studies for area 35 provided evidence for projections, and all studies on area 36 showed consistent projections only to the ventral striatum, (Table 2b), except for one lesion study showing associative projections from area 35 [153]. One study on the cingulate area 24a provided evidence of projections to the medial, central, and ventral striatum [98].

#### 3.1.3 Associative regions

For areas 18, 19, 23, 37, 38, 44, 45 and 46, several studies provide evidence for projections to the central and medial striatum only (Table 2c). For area 8, most studies (93.33%) found projections to the central and rostral striatum (Table 2c); except for one older lesion study, which suggested the presence of both dorsolateral and central striatal projections [70].

Areas 9 and 10 project only to central striatal regions (Table 2c), except one study showing ventral and central striatal projections [88]; however, this was a retrograde study where the striatal injection field was not specifically described. Most studies (77.78%) described Area 21 as having central projections, with two studies (22.22%) noting ventral striatal projections (Table 2c). Central striatal projections were only found in most studies that examined area 22 (73.33%) (Table 2c).

#### 3.1.4 Limbic and associative regions

Most studies on areas 25, 26, 31, 32 and 43 found consistent evidence for ventral and central striatal projections (Table 2d). Eight studies (66.67% of all included studies for area 11) found evidence only for central projections from area 11, however, four studies (33.33% of all included studies for area 11) consistently noted the presence of both central and ventral striatal projections (Table 2d). Based on these data we therefore considered area 11 to be projecting to both. The same logic applies to areas 12, 13, 14, 20 and 24b (Table 2d). These cortical areas were considered limbic and associative, based on projections to the ventral, dorsal and central striatum.

#### 3.1.5 Sensorimotor and limbic regions

Two studies provided evidence for consistent projections to the ventral limbic and dorsolateral sensorimotor striatum of areas 15 and 16 [95, 132] (Table 2e).

#### 3.1.6 Sensorimotor and associative regions

All studies that described striatal projections from areas 5 and 39 provided evidence of projections to both dorsolateral and central regions (Table 2e).

For area 40, four studies (66.67%) provided evidence for central projections while two studies (33.33%) provided evidence for projections to both dorsolateral and central striatal regions (Table 2e).

Six studies (60%) of the lateral area 7 (7l) provided evidence for central projections, and four studies (40%) provided evidence for projections to dorsolateral and central regions (Table 2e). Since both dorsolateral and central projections have been found in anterograde and retrograde studies, areas 40 and 7l were classified as both sensorimotor and associative (Table 2e).

#### 3.1.7 Regions without consistent striatal projections

Striatal projections from area 17, the primary visual cortex, have not been consistently demonstrated [84, 123], although older lesion studies did suggest the presence of connections to both the dorsolateral and central striatum [84, 96].

From areas 41 and 42, the primary auditory cortex (Table 2e), some studies demonstrated limited projections to the putamen and caudate [111, 151], while other studies have failed to demonstrate corticostriatal projections from these areas [161]. Due to a lack of consistent evidence of projections, these were considered as areas without striatal projections (Table 2e).

#### 3.2 Creation of the corticostriatal projection (CSP) atlas

Individual labels of the atlas were classified according to the overview provided in the Tables 2a-e. Since area 7m and 7l were assessed to likely have divergent projections to the striatum also in humans, we subdivided the PALS-B12 label for this area into a medial and lateral part. The results of this labelling are shown in Figure 2.

**Figure 2.**
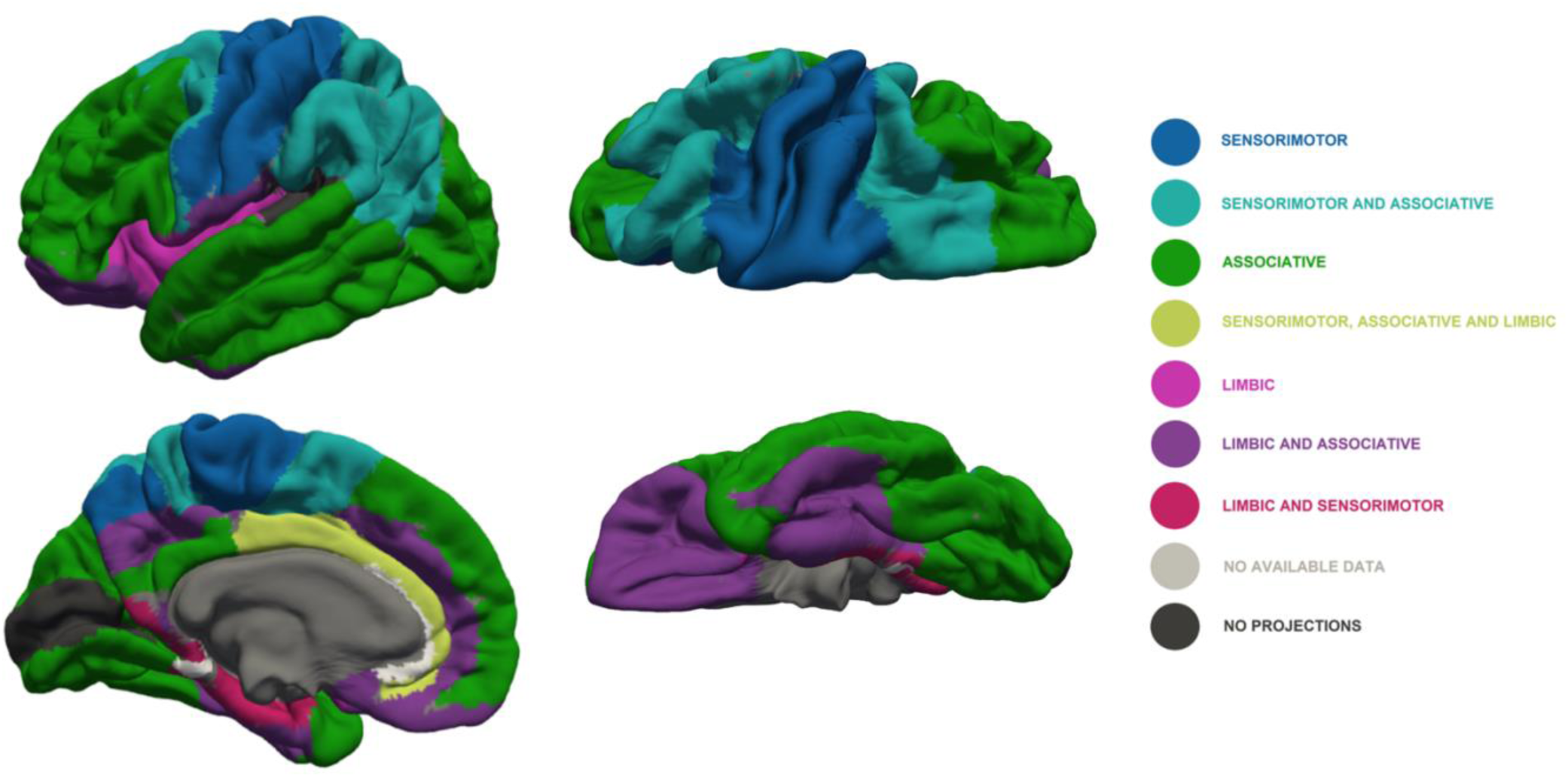
Lateral (upper right), medial (lower right), dorsal (upper left), and ventral (lower left) views of cortical areas, colors denote projection to terminal fields within different striatal subregions according to legend (left). *Note: Colors should be used for all figures in print*.

#### 3.3 DWI-based probabilistic tractography analysis

The results from the diffusion-weighted tractography analyses are shown in Figure 3. The connectivity density based on the number of streamlines was highest between the limbic atlas labels and the ventral striatum.

**Figure 3.**
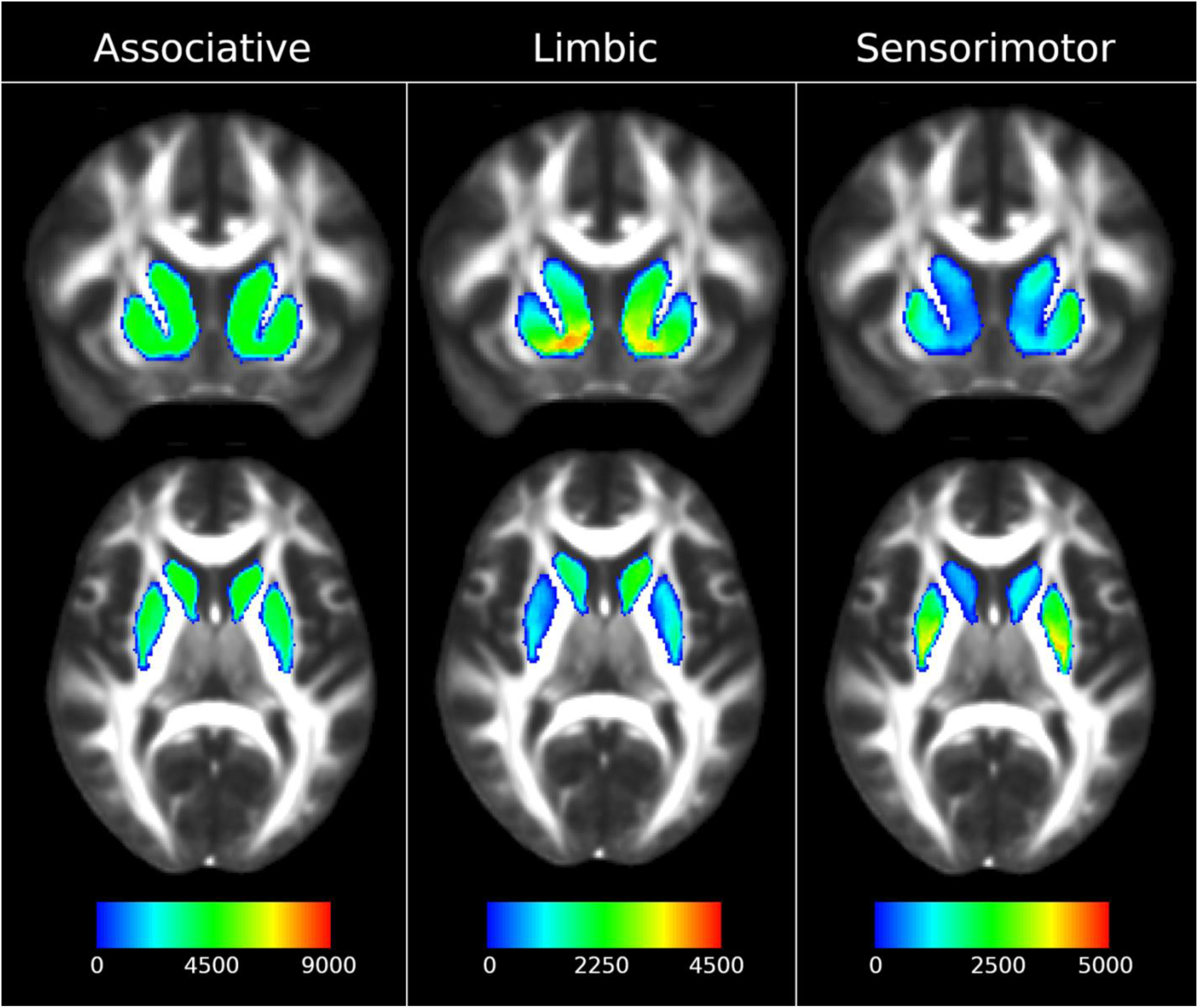
DWI analysis results of sensorimotor, associative, and limbic striatal target areas without masks for overlapping target areas. The color map indicates the number of valid streamlines originating in seed voxels in the striatum mask that terminate in the target cortical region. Scale bar: number of streamlines. *Note: Colors should be used for this figure in print*.

Similarly, for the sensorimotor atlas label, connectivity density was highest between the sensorimotor atlas labels and the dorsal and lateral aspects of the striatum. Connectivity density between the associative atlas label and the striatum was more diffusely spread throughout the dorsomedial striatum. The correspondences between these labels and connectivity densities were in line with our hypothesis with regards to the topography of the striatal target areas for cortical projections. We refer to Supplementary figures 1 and 2 for further details on the DWI analyses results.

## 4. Discussion

We present a new atlas of the human cortex according to striatal projections within different corticostriatal functional circuits, based on a comprehensive review of the primate anatomical tracing literature and the PALS-B12 atlas. Our labelling scheme will be a useful tool for neuroimaging studies focusing on specific subcircuits within the corticostriatal circuitry in the healthy and disordered brain.

For instance, new perspectives of mood disorders view the symptoms of emotional dysregulation to be expressions of disordered activity within large-scale brain networks [162, 163]. This paradigm requires a detailed understanding of the neuroanatomy underlying the different subcircuits involved in generating normal or symptomatic brain activity. Targeted intracranial brain stimulation of subcortical structures including the ventral striatum, has been shown as a promising treatment of severe depression [163, 164]. However, results from previous clinical trials have been inconsistent with regards to treatment effects [165]. Our anatomically informed overview and atlas of the corticostriatal circuitry, will facilitate the specific targeting and manipulation by pharmacological or deep brain stimulation of functional subcortical circuits in the context of neurological and psychiatric illness to address this issue.

We suggest that the CSP atlas will also be highly useful for the study of other brain disorders involving dysfunction within corticostriatal brain circuits, including schizophrenia and psychosis. Several morphological, histological, PET, probabilistic mapping and MRI studies provide evidence for specific gray matter loss in associative areas, notably the frontal, parietal, and temporal regions [166–172]. Few studies, however, have examined the specificity of these findings to the associative corticostriatal subcircuit, for which we now have provided an anatomy-based parcellation scheme. For instance, disturbances in the associative circuit may be preferentially linked to deficits in cognitive and executive function; abnormalities in the limbic circuit to disturbances in motivation, and abnormalities in the sensorimotor circuit to tactile hallucinations, delusions of control and motor symptoms. Our CSP atlas will enable testing of these specific hypotheses and facilitate the study of emotional disturbances with a greater anatomical and functional specificity. In turn this will provide new insights into symptom-brain circuit associations and the interactions between components within specific brain networks implicated in psychiatric disorders.

### The spatial distribution of cortical projection areas

Viewing the atlas as a whole, several observations are consistent with previous research on the topography of corticostriatal projections. First, (i) medial and ventral cortical areas project mainly to the limbic striatum; (ii) frontal, parietal, and temporal cortical areas project to the associative striatum, and (iii) caudal regions (comprising the motor cortices) project to the sensorimotor striatum. This topographical distribution is consistent with the conjecture that cortical areas demonstrating relative expansion during recent evolution tend to support associative functions, while the more evolutionarily conserved medial and ventral regions primarily serve functions related to motivation, reward, and visceral functions [173]. Second, the cortical regions that project to the associative striatum occupy the largest area relative to the whole-cortex area (40.1%), followed by sensorimotor (13.71%) and then limbic (2.27%) regions (Table 3). A corresponding relative difference in the proportion of corticostriatal projections seems to be preserved in striatal terminal fields [174], and a comparably greater proportion of the striatum is found to be associative in primates compared to rodents [16]. Third, there is a notable degree of overlap between functional domains of the striatum, illustrated by the number of cortical regions projecting to more than one striatal target zone, which is consistent with a convergence of striatal projections [174].

### Correspondence with *in vivo* DWI and functional imaging studies

In their landmark study, Draganski and colleagues used DTI to study the voxel-wise structural connectivity profiles of 23 cortical target regions using the thalamus and basal ganglia as seed regions [28]. They found evidence for a rostro-caudal gradient of frontal connectivity with the basal ganglia: The medial and orbital prefrontal cortex (MPFC and OFC) showed preferential structural connectivity with the ventral striatal region, the dorsolateral prefrontal (DLPFC) cortex projecting more diffusely to rostral and caudal aspects of both the caudate and putamen, the premotor (PM) cortex to dorsolateral putamen, and motor cortex (M1) to dorsolateral putamen and dorsal caudate. This pattern largely corresponds to our review of the primate anatomical literature. Specifically, our review suggests limbic or limbic/associative projections from cortical areas 10, 11 and 12, and the rostral part of area 32 (MPFC), and from areas 11, 12, 13 and 14 (OFC). For area 10, we found evidence for projections mainly to the associative regions and chose to classify this region as associative, although we note that additional limbic projections to the ventral region were found in some studies. Draganski and colleagues, however, chose to separate dorsal and ventral parts of area 10 into different labels. This may explain the discrepancy with our results, as dorsal area 10 mainly has associative projections, and ventral area 10 is mainly limbic. The choice to not delineate area 10 into dorsal and ventral parts was made after careful consideration of homology and corticostriatal targeting from primate anatomical data, since area 10 is cytoarchitectonally homogenous [175]. Therefore, a dorsal and ventral subdivision of area 10 would require an arbitrary delineation we could not support with our data. The DLPFC includes parts of areas 8, 9, 10 and 46, all predominantly associative areas, where findings were consistent between both our work and that of Draganski et al. The PM covers the entire area 6, M1 covers area 4 and for both these sensorimotor areas our results were again consistent with the findings of Draganski et al. By contrast, Draganski et al. did not identify connections from area 6 to the caudate, which is an established finding in primate studies. Notwithstanding these differences, the general pattern of topography found by Draganski et al. is consistent with our literature review.

In another prominent study, Tziortzi et al. (2014) combined DTI and PET and performed a functional analysis of dopamine release based on corticostriatal connectivity, dividing the cortex into occipital, temporal, parietal, and frontal lobes, and subdividing the frontal lobe further into caudal motor, rostral motor, executive and limbic subdivisions. Striatal terminal fields were then estimated as functional regions as a measure of regional dopamine transmission. The frontal lobe was found to dominate input to the striatum with about 82% of the striatal volume. The authors do not, however, report connection probabilities from the temporal and occipital lobes, as these regions were too small and spatially inconsistent across individuals. Furthermore, the study did not detect limbic connections from area 32/24, possibly due to the limited resolution of the imaging methods. It is also worth noting that this study found the highest degree of amphetamine-induced dopamine release in the limbic striatum, contrary to more recent studies [45]. The neuromodulatory effects of dopamine signaling from the striatum are shown to affect distal cortical regions [176–179]. These findings highlight the key role of the corticostriatal circuitry in the regulation of functional brain networks. In this context, the CSP atlas, as the first anatomically based overview of corticostriatal connections, will be useful for further studies of the role of the striatum in coordinating large-scale brain network activity.

In summary, although our approach differed from prior studies as discussed above, notable features of our review-based labelling scheme from primate studies show correspondence with previous results from anatomy and DTI studies. It is important to note however that this comparison is limited by some factors: First, the studies discussed above focused primarily on the frontal lobe, making it more difficult to compare precisely with our labelling in other cortical areas. Second, given the methodological limitations of DTI, e.g., limited resolution and the inability to ascertain the directionality of connections, it is impossible to validate the precise boundaries between projection areas through this comparison.

### Validation using DWI-based probabilistic tractography

To test for preferential connectivity of our atlas on *in vivo* human brain imaging data, we used probabilistic tractography to map the connectivity between the striatum and our cortical labels. Estimated surface areas of the cortex according to corticostriatal target area corresponded with our findings from primate anatomical data (Supplementary table 1). Our study was focused on the creation and testing of the CSP atlas, investigations of pathological brain states were beyond the scope of our study. Therefore, only healthy human brain DWI-data was used to test our model, but it will be important to extend this work by assessing differences between brain states in health and disease with the CSP atlas, which we make freely available to the community to do so. For example, the characteristics of specific disturbances in the corticostriatal circuitry could be studied by assessing DWI metrics suggestive of white matter abnormalities [180]. Studying distinct parts of the corticostriatal circuitry could be used to assess whether pathology is localized to one part of the circuit, as well as monitor individual treatment response to target treatment, for instance using PET studies with dopamine D2 receptor ligands.

### Strengths and limitations

Our study is the first extensive systematic review of the literature documenting cortical projections to the striatum based on the anatomical tracing studies in non-human primates with close to whole-cortex coverage. We found a notable consistency in the description of projections and striatal terminal fields across different studies using various tracing methodologies, supporting the robustness of our findings.

A clear pattern of projections was identifiable for most cortical regions included in our literature review, and both non-overlapping and overlapping projections were defined. Still, the tripartite subdivisions of striatal domains used for the cortical labeling is a functional model, and not based on macroscopic anatomical boundaries, with gradients of connectivity in boundary areas [1]. Furthermore, it may be argued that the use of striatal domains to label cortical projections presents a problem of circularity, given that the functions of the cortical regions itself are what defines striatal domains in the first place. We propose, however, that the ability to label corresponding regions in the cortex and striatum, belonging to distinct functional circuits may be a useful tool in the context of MRI analysis, despite this lack of accurate boundaries at cortical and subcortical levels.

We focused on dense projections as evidenced by anatomical tract tracing, however, diffuse projections also contribute to the striatal terminal fields and are found across functional subdivisions in overlapping zones. Furthermore, no distinction was made between projections from cortical layers 3 vs. 5 or projections from neocortical intra-telencephalic vs. pyramidal tract neurons [181], nor between projections to striosome, and matrix compartments [182, 183]. It is likely that the projection pattern found through our review is primarily descriptive of projections from layer 5 to the matrix, as this is the dominant projection type [184].

The translation between results from non-human primate studies into a human brain atlas is not without challenges. We note that the assessment of homology between the cortical anatomy and function across different primate species is a complex and evolving field [185, 186]. Notably, cortical regions undergoing expansion in the human brain compared to other primates seem to be mostly located in associative areas. This may entail a higher degree of imprecision with regards to homology, compared to more preserved sensorimotor and limbic areas. We refer to Supplementary materials, section 2, for a detailed discussion of considerations pertaining to homology of cortical brain regions.

While we relied on the Brodmann cytoarchitectural classification in our study, we considered several human brain labelling schemes based upon cytoarchitecture or cortical architecture using landmarks such as cortical gyri and sulci. As no authors used stereotaxic coordinates to describe cortical areas, brain atlases based upon stereotaxic coordinates, e.g., the Paxinos atlas [187], were not considered. Similarly, as gyri and sulci in primates may show individual variation with limited author description and difficulty with homological translation of these, the Destrieux-Halgren parcellation scheme [188] was also considered not suitable for our purpose. Designating cortical regions according to Brodmann classification based on author descriptions and figures is challenging but was more tractable than the above approaches. Thus, our evaluation was that on balance, the PALS-B12 atlas represented a robust choice for classifying corticostriatal projections based on Brodmann areas in primates. We acknowledge that new findings regarding primate homology may alter the implication and application of our results and digital atlas.

### Conclusion

We performed a comprehensive systematic review of anatomical studies on corticostriatal projections in non-human primate brain studies and present the first anatomically informed human cortical atlas of corticostriatal projection areas. Using *in vivo* diffusion-weighted tractography in healthy human participants, we were able to show that cortical regions in this atlas show preferential connectivity to functional striatal subregions. This atlas will serve as an anatomically informed foundation for the study of the different functional corticostriatal circuits in both the healthy and disordered brain.

## Acknowledgements

We are grateful and indebted to the anatomists, scientists and researchers involved in the studies reviewed, who through their dedicated and thorough work have provided a solid framework for future investigations.

