## Supplementary materials for "A new cortical parcellation based on systematic review of primate anatomical tracing studies on corticostriatal projections"

#### 1 Diffusion weighted imaging validation of the cortico-striatal projection atlas

The goal of the diffusion weighted imaging (DWI) validation was to test if the cortical regions of the CSP atlas showed the expected preferential connections to the striatum, as measured using structural connectivity. DWI preprocessing and probabilistic tractography was performed on the TSD (Service for Sensitive Data) computing cluster at the University of Oslo.

#### 1.1 DWI data acquisition

We included 24 healthy participants (58.3% female, mean age = 35.3, range = [20.1 – 51.5]) with no history of brain injury or neurological and neuropsychiatric disorders thought to affect brain structure. DWI was performed at Oslo University Hospital on a GE Discovery MR750 3T scanner with 32-channel head coil. The sequence parameters were: TE/TR = 81.3/8150 ms, resolution 2 $mm^3$ isotropic, b = 0, 1000, 2000, 3000 $s/mm^2$ with 60 non-coplanar diffusion directions for each b-shell. In addition, seven b = 0 volumes with reversed phase-encoding direction were acquired.

#### 1.2 DWI preprocessing

DWI data was preprocessed using an optimized pipeline. Briefly this pipeline corrects the data for noise, Gibbs ringing, echo-planar imaging (EPI), motion, eddy current, and susceptibility distortions. For more details, see Maximov, Alnæs and Westlye (2019) [1]. Fractional anisotropy (FA) maps [2] were estimated for each participant using a diffusion kurtosis approach [3]. Non-linear transformations between diffusion and MNI [4] spaces were computed with the *fsl_reg* command with FA maps as inputs.

#### 1.3 Selection of seed and target masks

As seed, we used a mask of the entire striatum, which was taken from the Oxford-GSK-Imanova structural atlas [5]. As targets, we used two sets of cortical masks. The first set contained the hub regions, BA 4, BA 9-10, and BA 35-36 and this analysis was used as a validation of the approach. The second set contained all the cortical regions of the CSP atlas, where cortical regions that project to more than one area of the striatum were represented by separate masks.

To create the target masks, we used the *mri_label2vol* command in FreeSurfer version 6.0.0 on the labels of the CSP atlas with the --*fill-ribbon* flag and a --*filthresh* of 0.5. To achieve an accurate transformation of the target masks from fsaverage to MNI space, we used the Registration Fusion (RF) approach [6] implemented in the Computational Brain Imaging Group (CBIG) toolbox [https://github.com/ThomasYeoLab/CBIG]. These masks were then mapped from MNI space to diffusion space using the transformations determined from the FA maps.

#### 1.4 Voxel-by-ROI connectivity

To estimate the connectivity from each voxel in the seed mask to each of the cortical target masks, we used the voxel-by-ROI connectivity approach implemented in FSL version 6.0.5 (FMRIB Software Library; [7]). First, a Bayesian estimation of diffusion parameters was obtained using sampling techniques for modeling crossing fibers (*bedpostx*) with the zeppelin deconvolution model (i.e. *-model* flag set to 3) [8, 9]. Briefly, the *bedpostx* command builds a distribution of diffusion parameters at each voxel and prepares the data for probabilistic tractography.

Voxel-by-ROI connectivity was computed using the *probtrackx2* command in FSL version 6.0.5 using the output from *bedpostx*. For the *probtrackx2* command, we set targets as waypoints with --*waycond=OR*. All target masks were merged to form a single stop mask. We enabled random samplings of points within a sphere with a radius of 1 centered on each voxel in the seed mask using the --*sampvox* flag. We also enabled random sampling of initial fibers using the --*randfib* flag. To improve the accuracy of the probabilistic tractography, we used the –*loopcheck*, --*onewaycondition*, and --*modeuler* flags. Seeds-to-target output was specified with the --*os2t* flag.

#### 1.5 Visualization of voxel-by-ROI connectivity results

Voxel-by-ROI connectivity maps and streamlines were inspected visually and transformed from individual diffusion space to MNI space. We then created group averages in MNI space for both voxel-by-ROI connectivity maps and streamlines. Figures were created showing the relative strength of the connectivity of each voxel in the striatal seed mask with each of the cortical target masks. To illustrate the valid streamlines, we also created figures showing valid streamlines for each of the cortical target masks.

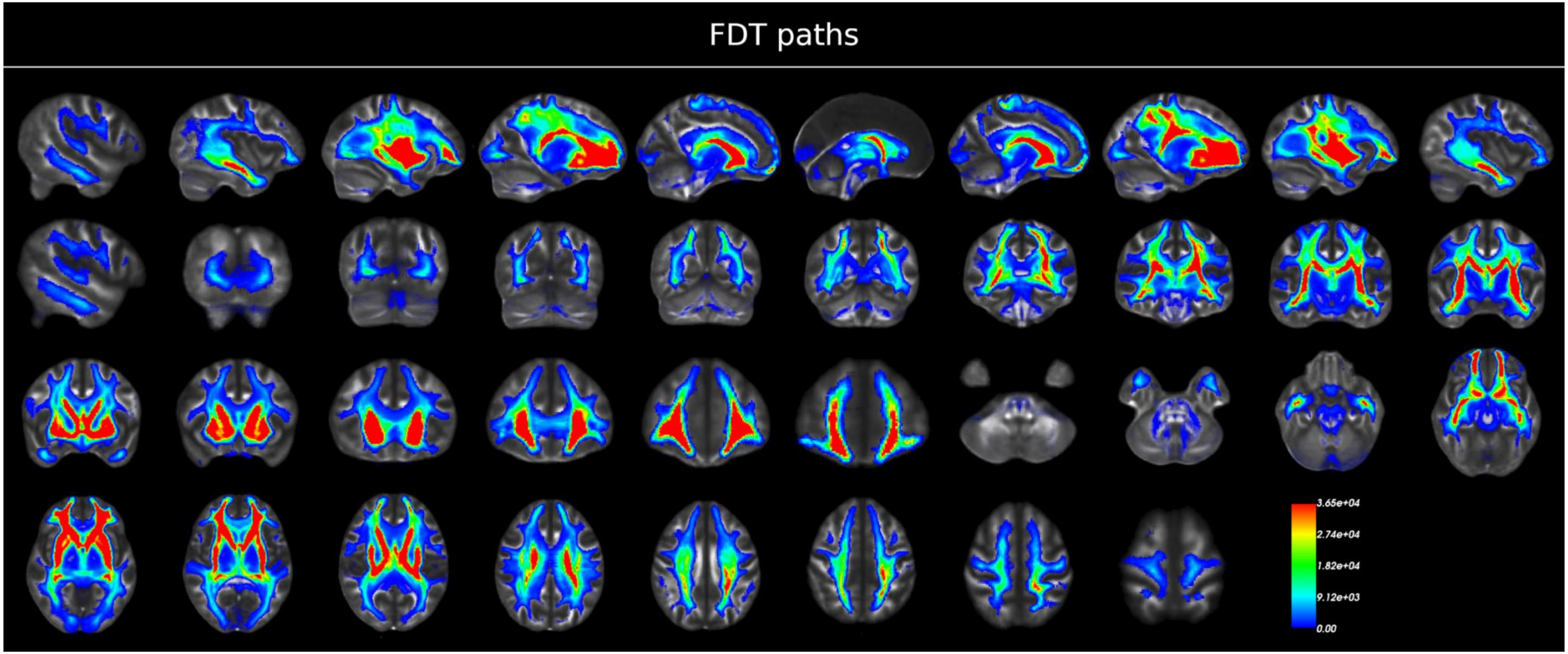


**Supplementary figure 1. DWI analysis results for fdt_paths tractography analysis**. *Note: Colors should be used for this figure in print.*

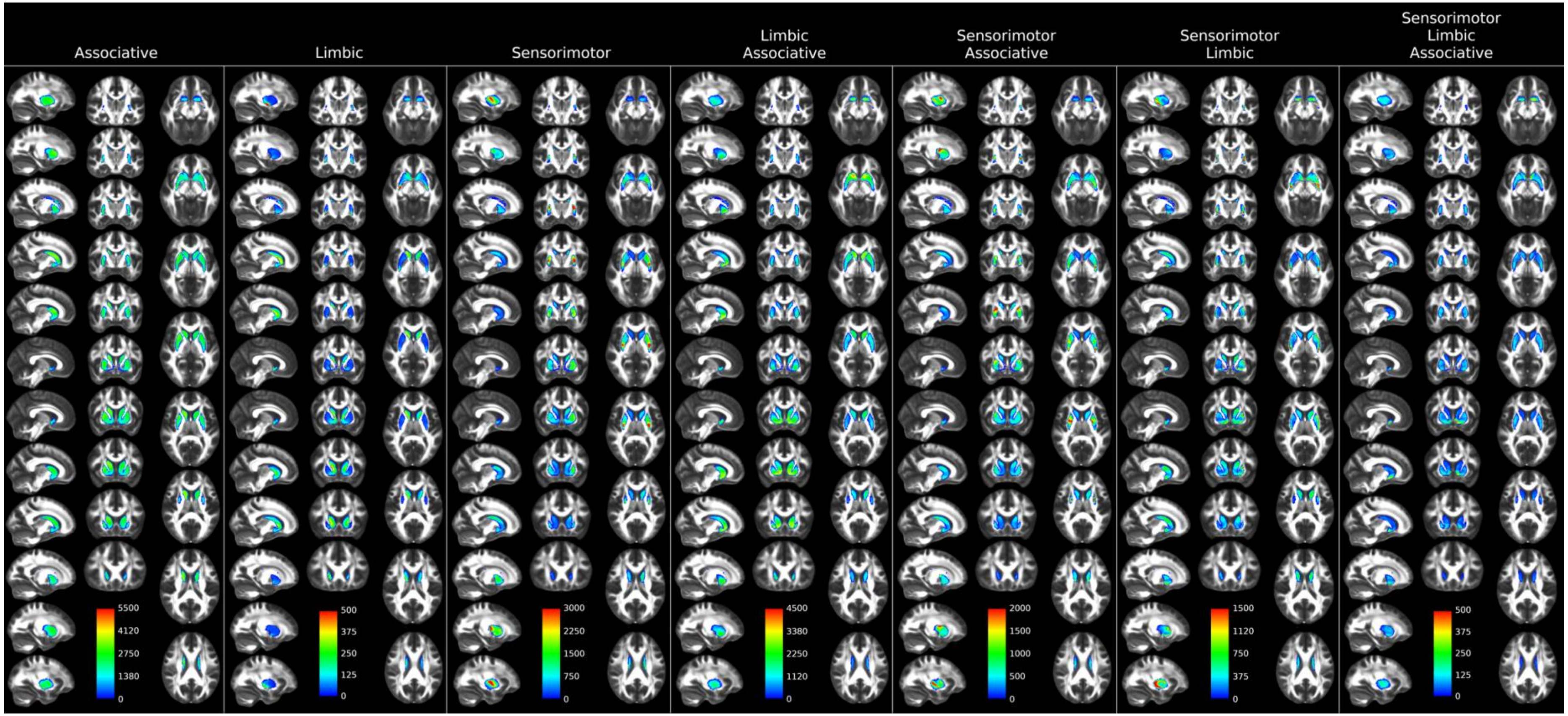


**Supplementary figure 2.** DWI analysis results for seeds-to-target tractography analysis, classified according to striatal target zones. *Note: Colors should be used for this figure in print.*

**Supplementary table 1. Estimated surface areas of the cortex according to corticostriatal target area.**

| ROI | Mean Area ($mm^2$) | Mean Area (%) |
|---|---|---|
| **Associative** | 29,891.90 | 40.10 |
| **Sensorimotor** | 10,223.47 | 13.71 |
| **Limbic** | 1,689.23 | 2.27 |
| **Limbic/Associative** | 9,867.15 | 13.24 |
| **Sensorimotor/Limbic** | 5,075.11 | 6.81 |
| **Sensorimotor/Associative** | 11,468.58 | 15.38 |
| **Sensorimotor/Limbic/Associative** | 1,955.20 | 2.62 |
| **No information** | 397.77 | 0.53 |
| **No projection** | 3,736.12 | 5.01 |
| **Whole cortex** | 74,551.68 | 100.00 |

### 2 Assessment of homologue cortical regions in the human brain

Establishing homology between cortical regions of humans and other non-human primates is a challenging task and an ongoing field of study. Human brains have undergone considerable growth in brain volume as well as increased gyrification of the cortical surface, resulting in a larger area and more complex configuration of the cerebral cortex compared to other primates. The evidence for homology between the human and non-human primate brain differs between cortical regions, and it has even been suggested that only homologies of visual cortex area 1 (V1), visual cortex area 2 (V2), middle temporal cortex MT and visual cortex area 5 (V5) are completely without dispute [10]. Yet, several anatomical aspects such as cytoarchitecture, topology, and preservation across non-human primate species as well as function may still be of use to determine an extent of homology. It is important that inquiries about homology between the human and non-human primate brain are defined specifically according to which of these features are considered.

Here, we consulted several reviews and book chapters to assess both cytoarchitectural and functional homology between the cortical regions investigated in several species of non-human primates and the corresponding Brodmann area of the human brain [10-18]. A supplementary table of all cortical regions labelled in the PALS-B12 atlas summarizes our findings in this article, including all included Brodmann areas with relevance to comparative anatomical studies. Another challenge encountered by using Brodmann areas, is that several sources disagree on the amount of Brodmann areas in both human and non-human primate brains [19]. We tried to mitigate some of this uncertainty by using cytoarchitectonic features as the primary source, supplemented with functional data for specific cortical regions where this was not available. For detailed considerations of homology, we refer to Supplementary table 2.

#### 2.1 Differences in sensorimotor and related areas in humans and non-human primates

Area 8 in the non-human primate brain has an established homology in the human brain with area 6. In the human brain, this area encompasses the frontal eye fields, while these are found in area 8 in non-human primates [20]. Due to this and some studies which suggest associative projections from area 6 may also exist in non-human primates, we chose to classify area 6 in the human brain as sensorimotor-associative. In the human brain, area 8 has undergone expansion with no clear homologous regions in simians [21]. Given the adjacency to area 9 we considered this to be best classified as an associative region.

Cingulate cortex area 24 is subdivided into three subregions [22]; areas 24a, 24b and 24c. In the macaque, areas 24a and 24b have both associative and limbic projections, whereas area 24c project to the sensorimotor striatum. In the human brain, this subdivision is also found, however, it is not clarified whether striatal projections follow a similar topography. Due to limitations of spatial resolution, it is not possible to accurately subdivide the PALS-B12 atlas into these subregions [22]. We therefore decided to denote area 24 in the human brain as collectively having projections to all functional domains.

Area 7 in the primate brain is often divided into its medial (7m) and lateral (7l) aspects. Here, an area expansion is found in the human brain, such that this area is homologous to regions 7, 39 and 40 in the human cortex [23]. Homology of the angular gyrus region (area 39) has been debated, with some researchers suggesting no clear homologous region exists in non-human primates [24], while others suggest areas 7a/PG exist in the macaque [25]. Area 40 has been described as a human-specific region that evolved with the development of tool-using skills [16]. Although it may not be considered known, this expansion likely originated from the lateral aspect of area 7. We therefore considered areas 7l, 39 and 40 in the human brain as homologous to primate area 7l and thus classified these regions as sensorimotor and associative.

#### 2.2 Differences in limbic and related areas in humans and non-human primates

There is a suggested functional homology between the macaque and human perirhinal area (areas 35 and 36), although with increased size and functional repertoire in humans [26]. Entorhinal areas are considered largely homologous between primate species [27, 28]. These areas are hence considered as limbic in the human brain.

Areas 15 and 16 of the insula in the temporal lobe are described as homologous to human areas 15 and 16, although Brodmann could not find this homology [29, 30]. These areas are considered sensorimotor and limbic, in line with tracing studies and their known functions in goal-directed reward behavior [31].

#### 2.3 Differences in associative and related areas in humans and non-human primates

Expansion and subdivision of non-human primate cortical areas is especially marked in prefrontal areas: guenon area 9 is homologous to human areas 9 and 10 [32], and guenon area 12 shows homology with human area 47/12 [33]. Macaque area 32 of the anterior cingulate gyrus is considered architectonically homologous to a subdivision of the human area 32, 32/prelimbic cortex (PL) [33]. The temporopolar area is comparable to human area 38, although it has undergone further expansion with increased heterogeneity in humans [34]. Macaque areas 44 and 45 are homologous to the same human areas [25, 35]. Hence, these areas were all classified as associative regions in the human brain.

Human dorsal area 31 (d31) in the cingulate cortex is described as homologous to macaque area 31 [36]. Guenon area 26 is considered homologous to human areas 26, 29 and 30 in the posteromedial or retrosplenial area [37]. These areas were considered as limbic and associative.

#### 2.4 Differences in areas without clearly established homology in humans and non-human primates

For certain areas specific to humans, no clear homologue region was found in the non-human primate brain. This includes human areas 33 (anterior cingulate cortex), 37 (fusiform gyrus), area 39 (angular gyrus), and area 40 (parietal region) [37]. For areas 37, 39 and 40, where the degrees of homology are debated, the pragmatic choice was made to include the comparable regions in non-human primate brains. For area 37, we considered face-selective regions active during face recognition as homologous to human area 37 [38]. In the primate area 37, only associative projections were demonstrated, and we considered this to be the best classification also in the human brain. Classification of projections from areas 39 and 40 are described above. Area 33 was excluded as no label exists for this region in the PALS-B12 atlas, and no studies were found describing relevant projections from here. Area 27 (piriform cortex) was excluded due to the lack of studies describing cortico-striatal projections from here.

**Supplementary table 2. Approximation of homology between human and primate Brodmann areas.**

| **Human Brodmann areas** | **Non-human primate homologue Brodmann area** | **Homologue species** | **References** | **Main differences between human and primate Brodmann areas** |
|---|---|---|---|---|
| 3,1 & 2 | 3, 1 & 2 | Guenon | [11, 32, 39] | |
| 4 | 4 | Guenon | [11, 32, 39] | BA4 occupies a much greater fraction of the monkey frontal lobe compared to humans [32]. |
| 5 | 5 | Guenon | [11, 32, 39] | |
| 6 | 6 and 8 | Guenon | [11, 20, 32, 39] | Human area 6 is larger than area 4, however the monkey area 4 is larger than area 6 [32]. Area 6 in humans also contain frontal eyefields (FEF), however in monkeys FEF are located in BA 8 [20]. |
| 7 | 7 | Guenon | [11, 32, 39] | BA7 in monkeys is considered homologous to areas 7, 39 and 40 in humans, as only undifferentiated precursor zones for parietal areas exist in the guenon. |
| 8 | No clear homologue | N/A | [20, 21, 32] | |
| 9 | 9 | Guenon | [11, 32, 39] | Also homologous to human area 10, according to Brodmann 1909. |
| 10 | 9 and 10 | Guenon | [11, 39] [20, 32, 33] | Also homologous to human area 9, according to Brodmann 1909. Humans have a larger area 10 than monkeys with specialized cortical layers involved in connections with other higher-order association areas [20]. |
| 11 | 11 and 12 | Macaque | [33, 40] | BA12 in monkeys is considered homologous to BA11 in humans [40]. |
| 12 | 12 | Macaque | [33] | BA47/12 in humans is considered homologous to BA12 in monkeys [33]. |
| 13 | 13 | Macaque | [33] | |
| 14 | 14 | Macaque | [33] | |
| 15 | 15 | Macaque | [30, 32] | Homology not found by Brodmann, according to Brodmann 1909. |
| 16 | 16 | Macaque | [30, 32] | Homology not found by Brodmann, according to Brodmann 1909. |
| 17 | 17 | Several species | [10, 32] | |
| 18 | 18 | Several species | [10, 32] | |
| 19 | 19 | Several species | [10, 32] | |
| 20 | 20 | Guenon and macaque | [11, 32, 39, 41] | |
| 21 | 21 | Guenon and macaque | [11, 32, 39, 41] | |

| | | | | |
|---|---|---|---|---|
| 22 | 22 | Guenon and macaque | [11, 32, 39, 41] | |
| 23 | 23 | Guenon and macaque | [11, 32, 39] | |
| 24 | 24 | Guenon and macaque | [11, 32, 39] | |
| 25 | 25 | Guenon and macaque | [11, 32, 39] | |
| 26 | 26 | Guenon and macaque | [11, 32, 39] | BA26 in monkeys is considered homologous to BA26, BA29 and BA30 in humans [32]. |
| 27 | 27 | Guenon and macaque | [11, 32, 39, 41] | |
| 28 | 28 | Guenon and macaque | [11, 27, 28, 32, 39, 41] | BA28 and 34 in monkeys are considered homologous to the human entorhinal area [28]. |
| 29 | 26 and 29 | Guenon and macaque | [32, 42] | BA26 in monkeys is considered homologous to BA26, BA29 and BA30 in humans [32]. |
| 30 | 26 and 30 | Guenon and macaque | [32, 42] | BA26 in monkeys is considered homologous to BA26, BA29 and BA30 in humans [32]. |
| 31 | 31 | Macaque | [36] | Macaque BA31 is considered homologous to area d31 in humans, as there is no area v31 in macaques [36]. |
| 32 | 32 | Macaque | [32, 33] | BA32 in the macaque is considered architectonally equivalent to human 32pl and topologically equivalent to human 32ac. |
| 33 | No homologue region | N/A | [32] | Only found in humans [32]. |
| 34 | 34 | Guenon and macaque | [11, 27, 28, 32, 39, 41] | BA28 and 34 in monkeys are considered homologous to the human entorhinal area [28]. |
| 35 | 35 | Macaque | [26, 32] | There is a suggested functional homology between macaque and human perirhinal area, although its size and repertoire increased in humans [26]. |
| 36 | 36 | Macaque | [11, 32, 39] | Not sufficiently differentiated in guenons, but observed in the marmoset by Brodmann [32]. |
| 37 | Face-selective regions | Macaque | [38] | Face-selective regions in monkeys are considered homologous to BA37 in humans, however there is no real cytoarchitectural homology [38]. |
| 38 | Temporopolar areas | Several species | [32, 34] | Temporopolar areas in monkeys are considered homologous to BA38 in humans, however these regions have undergone further evolution and expansion in humans [34]. |
| 39 | 7a/PG or no homologue region | Macaque | [15, 24, 35, 43, 44] | The homology of BA39 or the angular gyrus in humans to monkey brain regions is debated. Some authors suggest no clear homologue [24, 43, 44], while others suggest homology between human area 39 and areas 7a/PG in the macaque [15, 35]. |

| | | | | |
|---|---|---|---|---|
| 40 | No homologue region | N/A | [16] | Possibly specific human region for tool-using according to one functional study [16]. |
| 41 | Auditory cortex | Macaque | [11, 32, 39, 45] | Not described in the guenon by Brodmann [11, 39], but considered homologous to auditory cortices in the human brain [45]. |
| 42 | Auditory cortex | Macaque | [11, 32, 39, 45] | Not described in the guenon by Brodmann [11, 39], but considered homologous to auditory cortices in the human brain [45]. |
| 43 | 43 | N/A | [32, 46] | |
| 44 | 44 | Macaque | [15, 25, 32] | This homologous region in macaque brains is smaller and located entirely in the caudal wall of the inferior ramus of the arcuate sulcus [25]. |
| 45 | 45 | Macaque | [15, 25, 32, 47, 48] | |
| 46 | 46 | Macaque | [11, 32, 39] | |
| 47 | 12 or 47 | Macaque | [11, 32, 39, 49] | |
| 48 | No homologue region | N/A | [11, 18, 32, 39] | Possibly human-specific region due to temporal lobe expansion during evolution [18]. |
| 49 | Parasubiculum | Macaque | [32] | |
| 50 | | N/A | [11, 19, 39] | Not sufficiently described by Brodmann [19]. |
| 51 | Prepiriform area | N/A | [11, 19, 39] | |
| 52 | Parainsular area or no homologue region | Macaque | [32, 50] | Possibly a human-specific region, but possible homologous to the parainsular area in macaques [50]. |

**3. Search strategy**

Search for: 4 and 10 and 18 and 29

Results: 464

Database: Ovid MEDLINE(R) ALL <1946 to September 10, 2019>
Search Strategy:
--------------------------------------------------------------------------------
1 exp Cercopithecidae/ (120563)
2 monkey.mp. [mp=title, abstract, original title, name of substance word, subject heading word, floating sub-heading
word, keyword heading word, organism supplementary concept word, protocol supplementary concept word, rare disease
supplementary concept word, unique identifier, synonyms] (51442)
3 non-human primate.mp. [mp=title, abstract, original title, name of substance word, subject heading word, floating
sub-heading word, keyword heading word, organism supplementary concept word, protocol supplementary concept word, rare
disease supplementary concept word, unique identifier, synonyms] (3857)
4 1 or 2 or 3 (144389)
5 exp Cerebral Cortex/ (348161)
6 cortex.mp. [mp=title, abstract, original title, name of substance word, subject heading word, floating sub-heading
word, keyword heading word, organism supplementary concept word, protocol supplementary concept word, rare disease
supplementary concept word, unique identifier, synonyms] (455763)
7 exp Limbic Lobe/ah, cy [Anatomy & Histology, Cytology] (2059)
8 exp Parahippocampal Gyrus/ah, cy [Anatomy & Histology, Cytology] (1025)
9 insula.mp. [mp=title, abstract, original title, name of substance word, subject heading word, floating sub-heading
word, keyword heading word, organism supplementary concept word, protocol supplementary concept word, rare disease
supplementary concept word, unique identifier, synonyms] (10348)
10 5 or 6 or 7 or 8 or 9 (577525)
11 Basal Ganglia/ (12788)
12 exp Corpus Striatum/ (62139)
13 striatum.mp. [mp=title, abstract, original title, name of substance word, subject heading word, floating
sub-heading word, keyword heading word, organism supplementary concept word, protocol supplementary concept word, rare
disease supplementary concept word, unique identifier, synonyms] (63081)
14 striatal.mp. [mp=title, abstract, original title, name of substance word, subject heading word, floating
sub-heading word, keyword heading word, organism supplementary concept word, protocol supplementary concept word, rare
disease supplementary concept word, unique identifier, synonyms] (34858)
15 caudate.mp. [mp=title, abstract, original title, name of substance word, subject heading word, floating
sub-heading word, keyword heading word, organism supplementary concept word, protocol

supplementary concept word, rare
disease supplementary concept word, unique identifier, synonyms] (25339)
16 putamen.mp. [mp=title, abstract, original title, name of substance word, subject heading word, floating
sub-heading word, keyword heading word, organism supplementary concept word, protocol supplementary concept word, rare
disease supplementary concept word, unique identifier, synonyms] (16043)
17 accumbens.mp. [mp=title, abstract, original title, name of substance word, subject heading word, floating
sub-heading word, keyword heading word, organism supplementary concept word, protocol supplementary concept word, rare
disease supplementary concept word, unique identifier, synonyms] (20311)
18 11 or 12 or 13 or 14 or 15 or 16 or 17 (122094)
19 exp Microscopy/ (540233)
20 exp Histology/ (362979)
21 Autoradiography/ (50687)
22 exp Horseradish Peroxidase/ (15783)
23 exp Neural Pathways/ah, cy [Anatomy & Histology, Cytology] (22245)
24 projections.mp. [mp=title, abstract, original title, name of substance word, subject heading word, floating
sub-heading word, keyword heading word, organism supplementary concept word, protocol supplementary concept word, rare
disease supplementary concept word, unique identifier, synonyms] (47282)
25 anterograde tracing.mp. [mp=title, abstract, original title, name of substance word, subject heading word,
floating sub-heading word, keyword heading word, organism supplementary concept word, protocol supplementary concept
word, rare disease supplementary concept word, unique identifier, synonyms] (812)
26 retrograde tracing.mp. [mp=title, abstract, original title, name of substance word, subject heading word,
floating sub-heading word, keyword heading word, organism supplementary concept word, protocol supplementary concept
word, rare disease supplementary concept word, unique identifier, synonyms] (2117)
27 anterograde labeling.mp. [mp=title, abstract, original title, name of substance word, subject heading word,
floating sub-heading word, keyword heading word, organism supplementary concept word, protocol supplementary concept
word, rare disease supplementary concept word, unique identifier, synonyms] (366)
28 retrograde labeling.mp. [mp=title, abstract, original title, name of substance word, subject heading word,
floating sub-heading word, keyword heading word, organism supplementary concept word, protocol supplementary concept
word, rare disease supplementary concept word, unique identifier, synonyms] (1668)
29 19 or 20 or 21 or 22 or 23 or 24 or 25 or 26 or 27 or 28 (957108)
30 4 and 10 and 18 and 29 (464)
